# SETDB1 modulates the TGFβ response in Duchenne muscular dystrophy myotubes

**DOI:** 10.1101/2023.06.28.546840

**Authors:** Alice Granados, Maeva Zamperoni, Roberta Rapone, Maryline Moulin, Ekaterina Boyarchuk, Costas Bouyioukos, Laurence Del Maestro, Véronique Joliot, Elisa Negroni, Myriame Mohamed, Sandra Piquet, Anne Bigot, Fabien Le Grand, Sonia Albini, Slimane Ait-Si-Ali

**Author notes:** In memory of our collaborator and friend Armando Felsani.

## Abstract

Overactivation of the TGFβ signaling in Duchenne muscular dystrophy (DMD) is a major hallmark of disease progression, leading to fibrosis and muscle dysfunction. Here, we investigated the role of SETDB1, a histone lysine methyltransferase involved in muscle differentiation. Our data show that, following TGFβ induction, SETDB1 accumulates in the nuclei of healthy myotubes, while being already present in the nuclei of DMD myotubes where TGFβ signaling is constitutively activated. Interestingly, transcriptomics revealed that depletion of SETDB1 in DMD myotubes leads to downregulation of TGFβ-target genes coding for secreted factors involved in extracellular matrix remodeling and inflammation. Consequently, SETDB1 silencing in DMD myotubes abrogates the deleterious effect of their secretome on myoblast differentiation by impairing myoblast pro-fibrotic response. Our findings indicate that SETDB1 potentiates the TGFβ-driven fibrotic response in DMD muscles, providing a new axis for therapeutic intervention.

**Key results:**

- TGFβ induces nuclear accumulation of SETDB1 in healthy myotubes
- SETDB1 is enriched in DMD myotube nuclei with intrinsic TGFβ pathway overactivation
- SETDB1 LOF in DMD myotubes attenuates TGFβ-induced pro-fibrotic response
- Secretome of TGFβ-treated DMD myotubes with SETDB1 LOF is less deleterious on myoblast differentiation

## INTRODUCTION

Histone lysine methylation/demethylation is a major epigenetic mechanism that controls chromatin states (*1*), with lysine 9 of histone 3 trimethylation (H3K9me3) being a hallmark of repressed chromatin (*2*). SETDB1 (SET Domain, Bifurcated 1, also called KMT1E, or ESET in the mouse) (*3*) is one of the H3K9 SUV39 KMT family. SETDB1 regulates many cellular states, such as stemness and terminal differentiation, including skeletal muscle terminal differentiation (*1, 3*). Although SETDB1 is mainly nuclear in proliferating myoblasts, where it represses terminal differentiation genes through H3K9me3 deposition, we have uncovered an original mechanism whereby murine Setdb1 is exported to the cytoplasm of differentiating myoblasts in a Wnt-mediated manner, allowing the de-repression of late differentiation genes and the formation of mature multinucleated myotubes (*4*). SETDB1 is associated with many human diseases including inflammatory bowel disease (IBD) (*5*), many cancer types (*6*), neuropsychiatric, genetic and cardiovascular diseases (*3*).

Dystrophinopathies are a spectrum of muscle genetic diseases caused by alterations in the *Dystrophin* gene, and Duchenne muscular dystrophy (DMD) is the most severe form, caused by total absence of the coded protein (*7, 8*). Dystrophin is essential for the maintenance of muscle membrane integrity by providing a strong mechanical link between the actin cytoskeleton and the extracellular matrix (ECM) (*9, 10*). Thus, the lack of functional Dystrophin is deleterious to membrane integrity, at the origin of the continuous degeneration of skeletal muscles (*11*), leading to exhaustion of muscle stem cells (MuSCs), progressive muscle wasting, chronic inflammation and severe fibrosis (i.e., accumulation of ECM components such as Collagens), which is the most prominent feature of DMD. Fibrosis is defined as pathological wound healing with an abnormal deposition of ECM resulting in the replacement of functional tissue by fibrotic connective tissue, which leads to an alteration of tissue and organ homeostasis (*12, 13*). Fibrosis involves intracellular and extracellular soluble effectors also called secretome, including pro- and anti-inflammatory cytokines, growth factors and ECM remodelers such as TGFβ, Matrix metalloproteinases (MMP)/Tissue Inhibitors of MMPs (TIMPs) and Collagens (*14*). The main pathway that sustains this excessive pro-fibrotic response is the TGFβ pathway, which is overactivated in DMD (*15, 16*).

In DMD patients, TGFβ levels, monitored by the nuclear accumulation of phosphorylated SMAD2/3 (pSMAD2/3) transcriptional modulators, are elevated in both plasma and muscle, and were shown to contribute to pathological fibrosis (*17*). However, the signaling pathways contributing to human DMD fibrosis, and the signals coordinating diverse cell types to maintain or restore muscle tissue homeostasis, remain largely unknown. Yet, TGFβ pathway overactivation is the common feature observed in Dystrophin-deficient myotubes *in vitro* independently of the type of mutation (*15, 16*) and exacerbated TGFβ/SMAD pathway has been shown to lead to fusion defects in *in vitro* DMD myotubes. Moreover, TGFβ pathway has been described as a molecular brake of myoblast fusion and, thus, muscle regeneration, while its inhibition leads to the formation of giant myofibers *in vivo* (*18*).

SETDB1 has been shown to regulate the TGFβ response in cancer and in T-cells (*19–22*) and pulmonary fibrosis (*23*) contexts. Here we investigated the SETDB1/TGFβ interplay in a model of DMD myotubes derived from hiPSCs or immortalized myoblasts (*15, 24*).

We show that SETDB1 contributes to the deregulated TGFβ signaling in DMD, indicating that SETDB1 may participate in muscle dysfunction and increased fibrosis. Indeed, TGFβ induces nuclear accumulation of SETDB1 in healthy myotubes, while SETDB1 is constantly accumulated in DMD myotube nuclei with intrinsic overactivated TGFβ pathway. Moreover, transcriptomics showed that SETDB1 silencing attenuates the TGFβ-dependent pro-fibrotic and anti-differentiation response in DMD myotubes. Interestingly, many TGFβ/SETDB1-dependent genes code for secreted proteins involved in ECM remodeling and inflammation. Conditioned medium assays show that TGFβ-treated myotubes with SETDB1 LOF produce a secretome which reduces the negative impact of TGFβ response on myoblast differentiation and impairs myoblast pro-fibrotic response. Our findings point to a role of SETDB1 in the fine-tuning of the TGFβ response in muscles, which is deregulated in DMD, participating in fibrosis.

## RESULTS

### SETDB1 relocalizes in the nuclei of healthy differentiated human myotubes in response to TGFβ pathway activation

All our experiments were conducted in human models generated from either human induced pluripotent stem cells (hiPSCs, which we have already described (*15*), or from immortalized myoblasts derived from healthy or DMD patients (*24*). hiPSCs can be directly induced to differentiate into myotubes using a transgene based epigenetic reprogramming strategy (*15*).

We have already showed that in murine myoblasts Setdb1 is both cytoplasmic and nuclear, and during terminal muscular differentiation Setdb1 is exported to cytoplasm in a Wnt-dependent manner, allowing de-repression of muscle genes (*4*). Thus, we have first investigated, and quantified (as depicted in **Figure S1A**), the subcellular localization of SETDB1 in proliferating *versus* differentiating healthy human myoblasts, and in myotubes in response to TGFβ/SMAD pathway activation (**Figure 1A**). TGFβ/SMAD pathway is constitutively activated in human proliferating myoblasts while nuclear phospho-SMAD3 (p-SMAD3) levels decrease upon differentiation (**Figures 1B, S1B**), as already shown in murine muscle cells (*18*). Differentiated myotubes are still responsive to TGFβ/SMAD pathway activation since p-SMAD3 nuclear levels are restored upon TGFβ1 treatment (**Figure 1F, S1B**). TGFβ treatment efficiency was evidenced by the activation of many TGFβ target genes, including *TGFβ* gene itself, concomitant to a decrease in many differentiation genes (**Figure S1D**).

**Figure 1:**
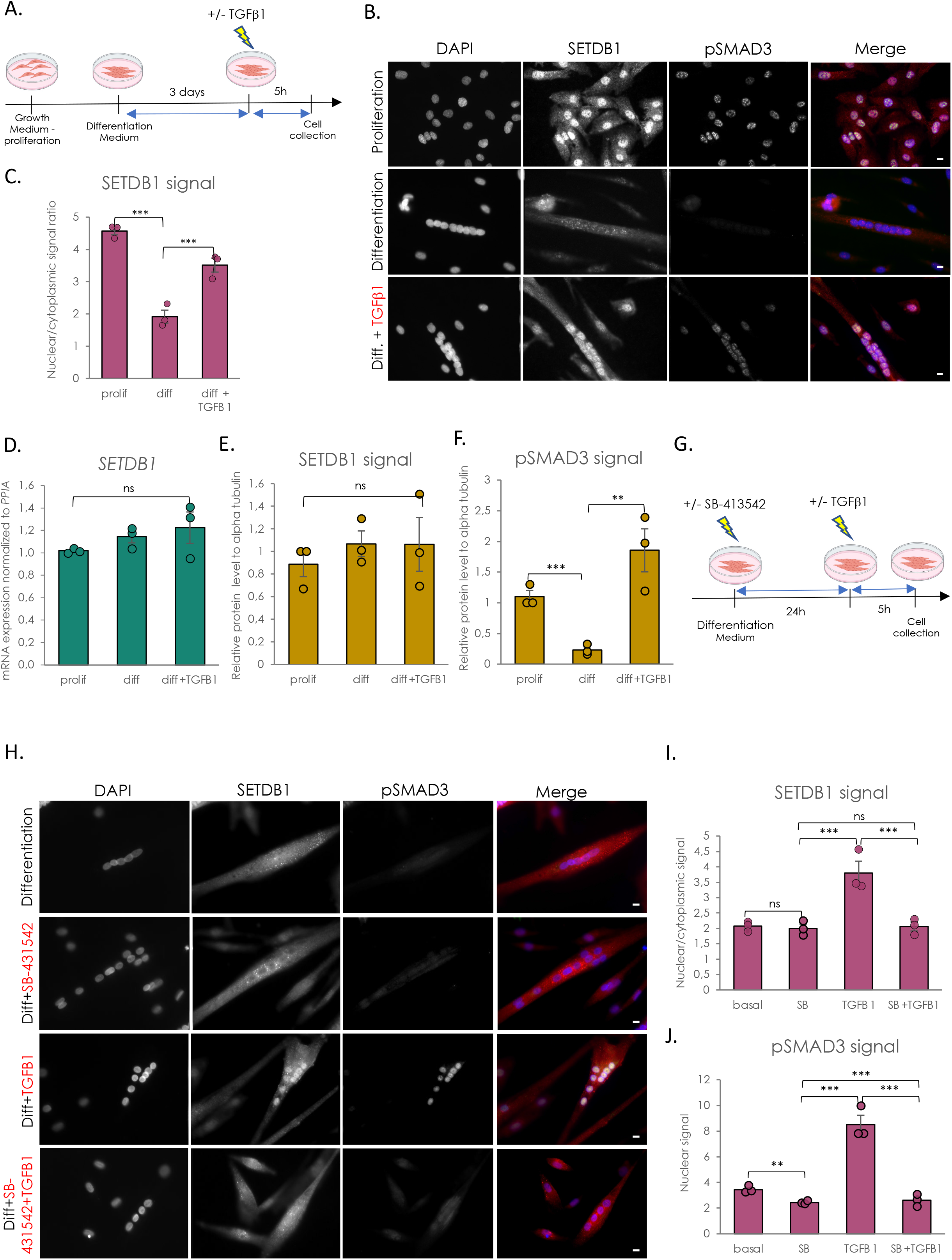
SETDB1 is excluded from myotube nuclei upon differentiation and can be relocated in response to TGFβ pathway activation. **A.** Scheme of experimental design. Cells were collected in proliferating phase (myoblasts) or after 3 days of differentiation (myotubes) with or without TGFβ1 treatment at 20 ng/mL. **B.** Immunostaining of SETDB1 (red) and phospho-SMAD3 (green). Nuclei were stained with DAPI (blue). Scale bar, 10 μM. **C.** Quantification of SETDB1 nuclear/cytoplasmic signal ratio. **D.** RT-qPCR of *SETDB1* in healthy proliferating myoblasts and differentiated myotubes +/- TGFβ1. **E-F.** Quantification of SETDB1 and phospho-SMAD3 protein levels in proliferating myoblast and myotube treated or not with TGFβ1 in total extracts. Alpha tubulin was used as a loading control to normalize samples. **G.** Scheme of experimental design with TGFβ inhibitor SB-413542. **H.** Immunostaining of SETDB1 (red) and phospho-SMAD3 (green). Nuclei were stained with DAPI (blue). SB-413542 blocks SETDB1 relocalization in myotube nuclei upon TGFβ1 treatment. Scale bar, 10 μM. **I-J.** Quantification of SETDB1 nuclear/cytoplasmic signal ratio (I), and phospho-SMAD3 nuclear signal (J). **For all panels:** Statistics were performed on ≥3 biological replicates (>100 nuclei for immunostaining quantification) and data are represented as average +/- SEM *p<0.05; **p<0.01; ***p<0.001 (unpaired Student’s t test).

We have confirmed in human muscle cells that SETDB1 is also highly enriched in the nuclei of proliferating myoblasts while it is depleted in nuclei of differentiated myotubes (**Figures 1B, 1C**), as in murine cells (*4*). Interestingly, TGFβ/SMADs pathway activation in muscle cells is correlated with SETDB1 presence in nuclei (**Figures 1B, 1C**) and we were able to trigger SETDB1 nuclear enrichment in differentiated myotubes after TGFβ1 treatment (**Figures 1B, 1C**), in absence of significant changes in SETDB1 mRNA (**Figure 1D**) and protein levels (**Figures 1E, S1C**), neither during terminal differentiation nor in response to 24h-TGFβ1 treatment. This result was confirmed in our healthy human hiPSCs-derived model (*15*) of muscle cells (**Figure S2C** upper panels). We next used the ALK4/5/7 inhibitor SB-431542 in myotubes to block TGFβ pathway activation prior to TGFβ1 addition (**Figure 1G**). The SB-431542 efficiency was evidenced by its effect on phospho-SMAD3 nuclear levels and TGFβ target and *MYOD1* gene expression (**Figures 1H, 1J, S1E**). Autocrine TGFβ signal blocking does not change SETDB1 localization in myotubes in basal condition but blocks SETDB1 nuclear relocalization in healthy myotubes in response to TGFβ1 (**Figures 1H, 1I**).

Altogether, these data showed that SETDB1 is excluded to the cytoplasm upon terminal differentiation of human myoblasts, and TGFβ pathway activation is directly responsible for SETDB1 nuclear relocalization in myotube nuclei.

### SETDB1 is accumulated in DMD myotube nuclei in TGFβ-dependent manner

We next investigated SETDB1 level and subcellular localization in the context of DMD in which the TGFβ pathway is overactivated both *in vivo* (*17, 25*) and in *in vitro* cellular models (*15, 16*).

First, immunohistochemistry assays show that SETDB1 co-localizes with phospho-SMAD3 in the centrally localized nuclei in DMD patient muscle histological sections (**Figure 2A**). To further confirm this observation in a more quantitative manner, we efficiently differentiated three healthy myoblasts or hiPSCs lines and three DMD lines carrying different *Dystrophin* mutations (myoblasts with exons 45-52 deletion (**Figure 2B**), myoblasts with point mutation in exon 59 (**Figure S2B**), and hiPSCs with exon 45 deletion (**Figure S2C**). Compared to healthy differentiated myotubes, DMD myotubes display overactivated TGFβ/SMAD pathway, as depicted by *TGFβ1* mRNA expression (**Figure S2A**) and nuclear phospho-SMAD3 protein level (**Figures 2D, 2F,** and (*15, 16*)), as expected.

**Figure 2:**
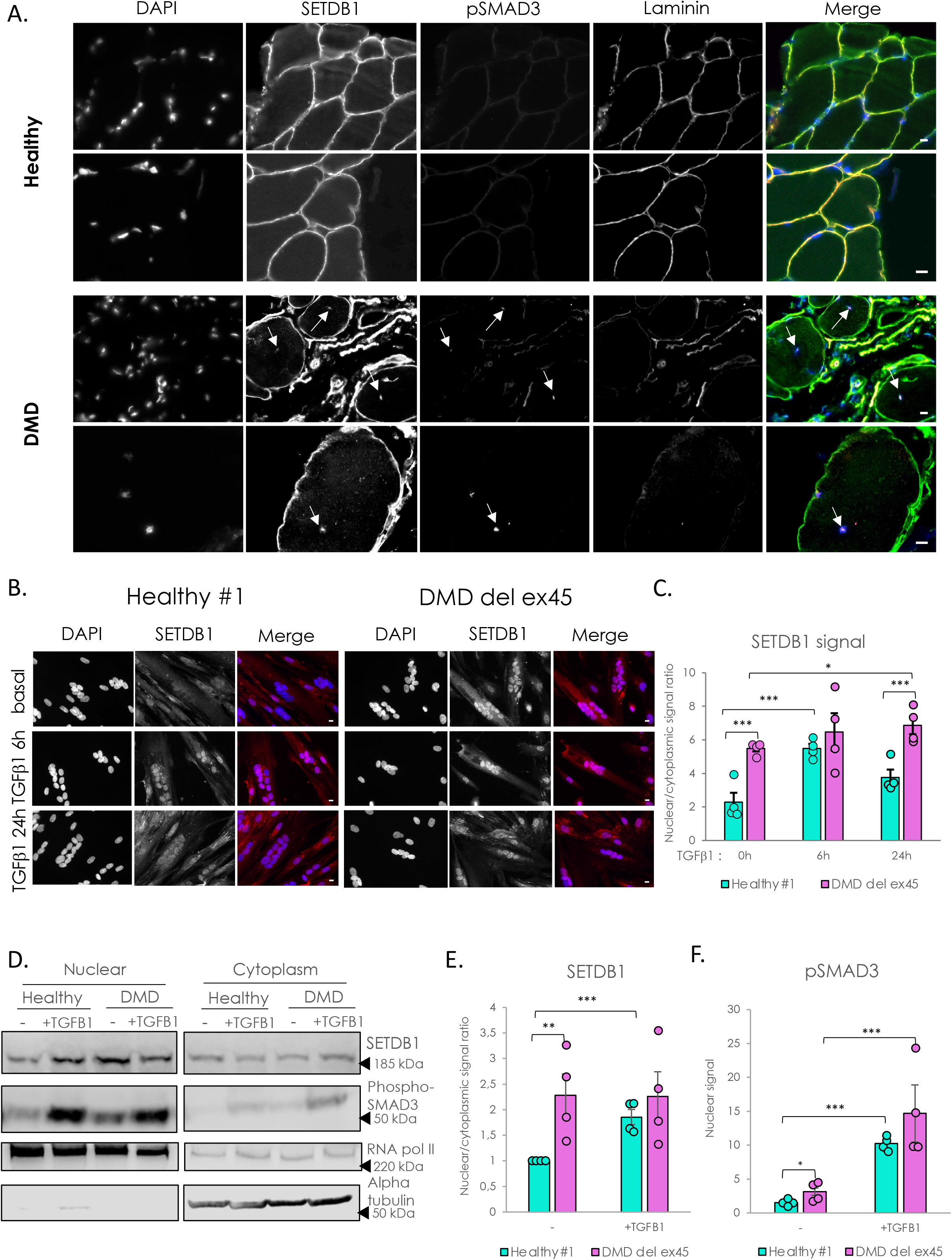
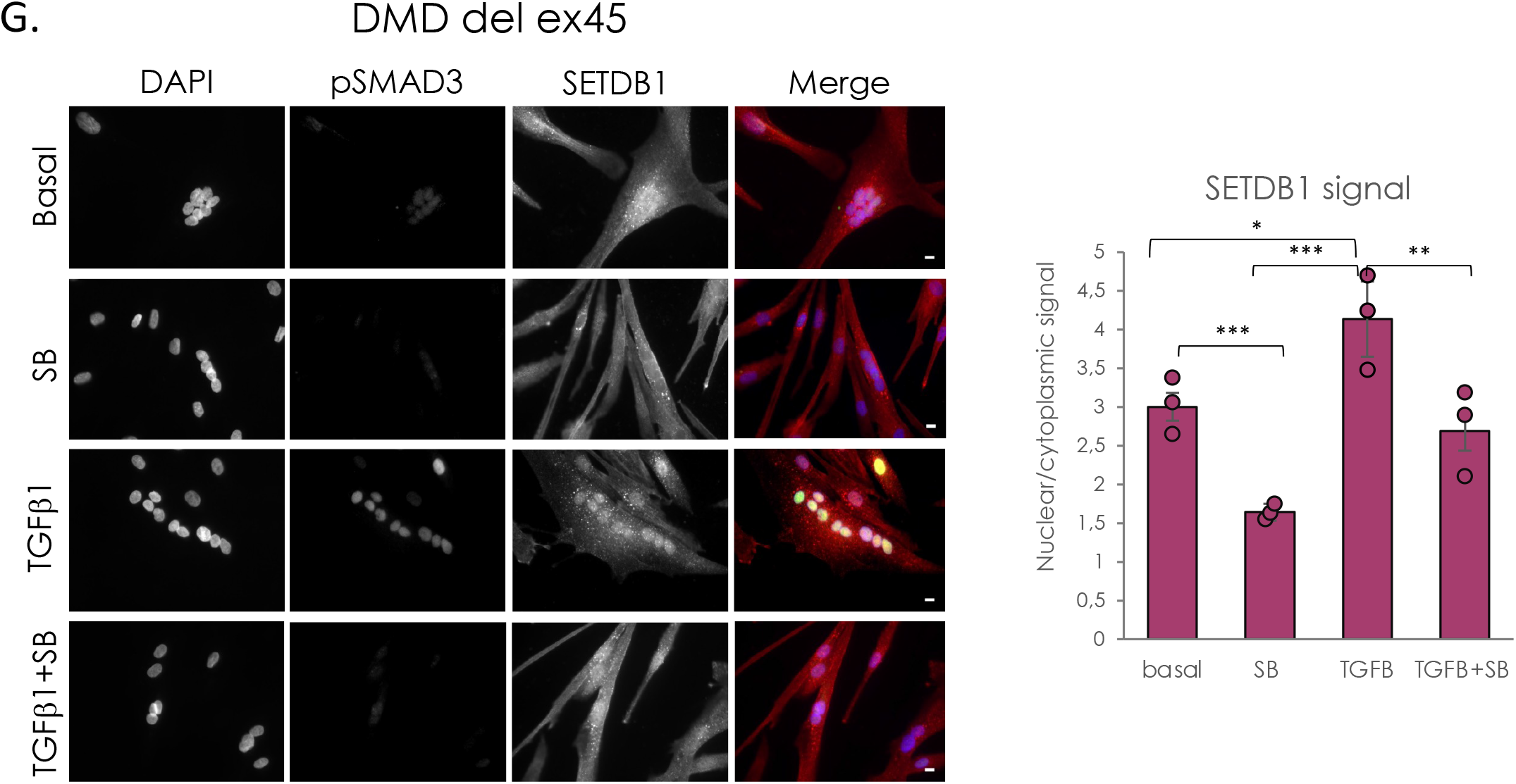
DMD differentiated muscle cells display a constitutive relocalization of SETDB1 in the nuclei and a higher activation of TGFβ/SMAD pathway. **A.** Immunostaining of SETDB1 (green), pSMAD3 (magenta) and laminin (red) on histological slides from healthy individual or DMD patient paravertebral muscles. Nuclei were stained with DAPI (blue). White arrows identify centrally localized nuclei of damaged myofibers in DMD muscle section. Scale bar, 10 μM. **B.** Immunostaining of SETDB1 (red) in healthy#1 and DMD del ex 45 myotubes. Nuclei were stained with DAPI (blue). Scale bar, 10 μM. **C.** Quantification of SETDB1 nuclear/cytoplasmic signal ratio. **D.** Western blot of nuclear and cytoplasmic fractions of healthy and DMD myotubes in response to TGFβ1 showing protein levels of SETDB1 and phospho-SMAD3. RNA polymerase II and alpha tubulin were used as loading controls of nuclear and cytoplasmic fractions respectively. **E-F.** Quantification of SETDB1 nuclear/cytoplasmic ratio (E), and phospho-SMAD3 nuclear signal (F). **G.** Immunostaining of SETDB1 (red) and phospho-SMAD3 (green) in DMD del ex45 myotubes and quantification of SETDB1 nuclear/cytoplasmic signal ratio. Nuclei were stained with DAPI (blue). Scale bar, 10 μM. **For all panels:** Statistics were performed on ≥3 biological replicates (>100 nuclei for immunostaining quantification) and data are represented as average +/- SEM *p<0.05; **p<0.01; ***p<0.001 (unpaired Student’s t test).

The quantification of SETDB1 nuclear/cytoplasmic ratio in healthy myotubes shows that SETDB1 translocated in the nuclei after 6h of TGFβ1 treatment and this relocalization tends to decrease after 24h (**Figure 2B-C, S2A-B**). Interestingly, in DMD myotubes, SETDB1 is accumulated in the nuclei at steady state and is persistent in response to TGFβ pathway activation (DMD exon 45 deletion in myoblasts or hiPSCs, **Figures 2B, 2C**; DMD point mutation, **Figure S2B**) and is associated with a higher activation of *TGFβ1* expression (**Figure S2A**). The effect of TGFβ pathway activation on the subcellular localization of SETDB1 is further confirmed by the western blotting of nuclear and cytoplasmic fractions of healthy and DMD myotubes (**Figures 2D, 2E, 2F**) and also in our hiPSC-derived DMD myotubes (exon 45 deletion, **Figure S2C**). We also blocked the autocrine signal of TGFβ in DMD myotubes with SB-431542 and found that it leads to decreased SETDB1 levels in nuclei with and without TGFβ1 treatment (**Figure 2G**). Overall, these data showed an effect of TGFβ on SETDB1 nuclear relocalization in at least 3 different DMD patient myotubes with different *Dystrophin* gene mutations.

To elucidate the molecular mechanisms underlying SETDB1 relocalization in DMD myotubes, we first tested if SETDB1 nuclear translocation in response to TGFβ was dependent on SMAD3, which is the effector translocating to the nucleus in response to TGFβ. Our results show that SETDB1 nuclear enrichment in response to TGFβ is not dependent on the presence of SMAD3 protein (**Figure S2D-F**).

SETDB1 is known to bear numerous post-translational modifications including many phosphorylations (https://www.phosphosite.org/proteinAction.action?id=9498&showAllSites=true). We thus hypothesized that the phosphorylation status of SETDB1 might correlate with its subcellular localization. To check this, we tested the migration profile of SETDB1 in several subcellular compartments. Our data showed that SETDB1 from the cytosolic and peri-nuclear fractions migrates higher compared to nuclear SETDB1 (**Figure S2G**), indicating possible hyperphosphorylation in these compartments. However, these observations require further investigation.

Taken together, these data showed that TGFβ induces SETDB1 nuclear enrichment in normal and DMD myotubes, regardless of the genetic background. This suggested that SETDB1 could modulate the cellular response to TGFβ.

### SETDB1 loss-of-function attenuates the TGFβ response in human DMD myotubes

To investigate the role of SETDB1 in the TGFβ response, we performed SETDB1 loss-of-function (LOF) in both healthy and DMD myotubes treated or not with TGFβ1 (as depicted in **Figure 3A**). While the siRNA mediated SETDB1 knockdown (KD, more than 80%) was efficient (**Figures 3B, S3A**), it did not show any effect on the levels of SMAD2/3 proteins nor their phosphorylation levels (**Figure 3B**). However, SETDB1 LOF led to a decrease in the TGFβ response, as depicted by the expression of *ad hoc* TGFβ target genes involved in pro-fibrotic response and inflammation, such as *TGFβ1*, *TIMP1* (*Tissue Inhibitor of Metalloproteinases 1*), *IL6* (*Interleukin 6*), *FN1* (*Fibronectin 1*) and *CTGF* (*Connective Tissue Growth Factor*) **(Figure 3C**, **S3C**). Note that SETDB1 KD alone, in absence of TGFβ, does not affect or only slightly the expression of these genes (**Figure S3D-E**). Since these TGFβ target genes are downregulated in SETDB1 KD condition, we hypothesized that in normal condition SETDB1 could target and repress inhibitors of the TGFβ pathway. A gene candidate approach showed that, indeed, SETDB1 LOF led to an increase in *SKIL* and *SMAD7* genes only in DMD (**Figures 3D, S3B**), two known inhibitors and fine tuners of the TGFβ pathway, while in absence of TGFβ, SETDB1 KD alone does not affect or only slightly the expression of these genes (**Figures S3D**, **S3F**). Of note, concomitant to the attenuation of the TGFβ response, SETDB1 LOF in 3-days differentiated myotubes induced a significant increase in muscle early differentiation marker *Myogenin* and regeneration marker *MYH3 (embryonic Myosin Heavy Chain*), but not in the muscle late differentiation marker *MYH1* (**Figures 3E, S3B**). Of note, SETDB1 KD alone also showed an effect on the expression of muscle differentiation genes (**Figure 3E, S3G**).

**Figure 3:**
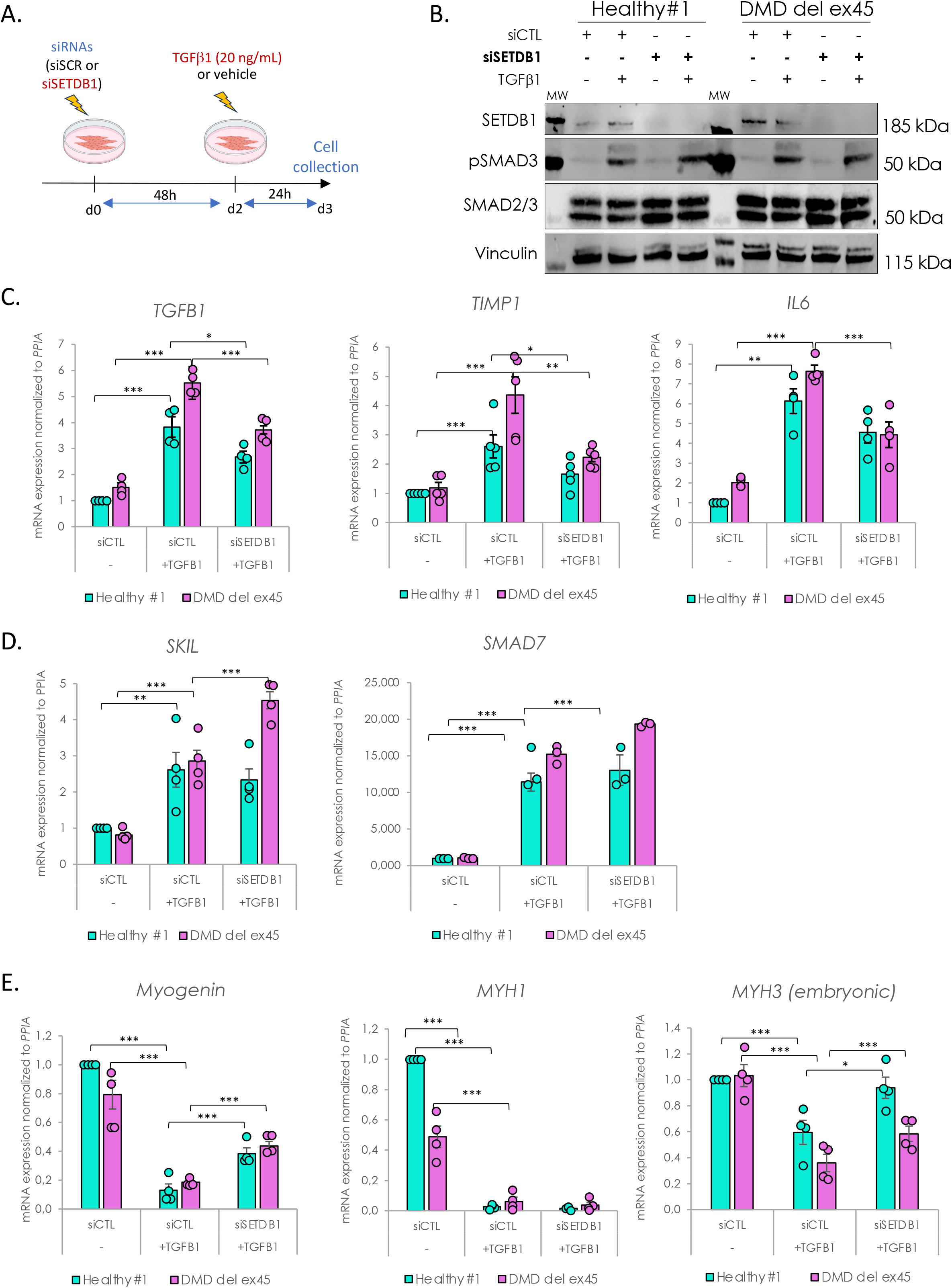
SETDB1 silencing leads to a decreased response to TGFβ1 in DMD myotubes while keys myogenic markers increase. **A.** Scheme of experimental design. Cells were treated with siRNAs scrambled (CTL) or against SETDB1 for 2 days after 3 days of differentiation (myotubes) and then treated or not with TGFβ1 at 20 ng/mL. **B.** Western blot showing SETDB1, phospho-SMAD3 and total SMAD3 protein levels. H3 was used as a loading control. **C.** RT-qPCR of TGFβ/SMADs pathway known targets *TGFβ1, IL6* and *TIMP1* in healthy and DMD myotubes +/- siSETDB1 +/- TGFβ1. **D.** RT-qPCR of TGFβ/SMADs pathway inhibitors *SKIL* and *SMAD7* in healthy and DMD myotubes +/- siSETDB1 +/- TGFβ1. **E.** RT-qPCR of myogenic markers *Myogenin, MYH1* and *MYH3 (embryonic MHC)*. **For all panels:** Statistics were performed on ≥3 biological replicates and data are represented as average +/- SEM *p<0.05; **p<0.01; ***p<0.001 (unpaired Student’s t test).

Thus, SETDB1 LOF attenuates the TGFβ1-induced fibrotic response while promoting regeneration and could be beneficial in DMD patients.

### SETDB1 alters the TGFβ regulated secretome in human DMD myotubes

To deepen the role of SETDB1 in the TGFβ response at the global level, we performed RNA-seq comparing DMD myotubes with healthy myotubes in response to TGFβ, with or without SETDB1 LOF. Principal component analysis (PCA) of gene expression levels showed that the triplicates and duplicates were distinctively separated and that the main source of variability between samples are the cell line source, healthy *versus* DMD. The second source seems to be the effect of the TGFβ treatment and SETDB1 silencing (**Figure S4A**).

To track changes in gene expression, first in response to TFGβ, we performed differential gene expression analysis (**Figure 4A**). We identified 1043 TFGβ-dependent differentially expressed genes (DEGs) in healthy myotubes (489 up- and 554 down-regulated genes) and only 480 in DMD myotubes (363 up- and 117 down-regulated genes) (Top genes in **Table S1, S2**). These data showed that while the TGFβ pathway is already intrinsically activated in DMD myotubes it is not saturated and can still be further activated *in vitro* (see also **Figure 2 & 3**, and (*15*)). Interestingly, only TFGβ-dependent 184 TFGβ-dependent genes (corresponding to 18% of healthy and 39% of DMD DEGs) are commonly deregulated in healthy and DMD conditions (**Figure 4A**).

**Figure 4:**
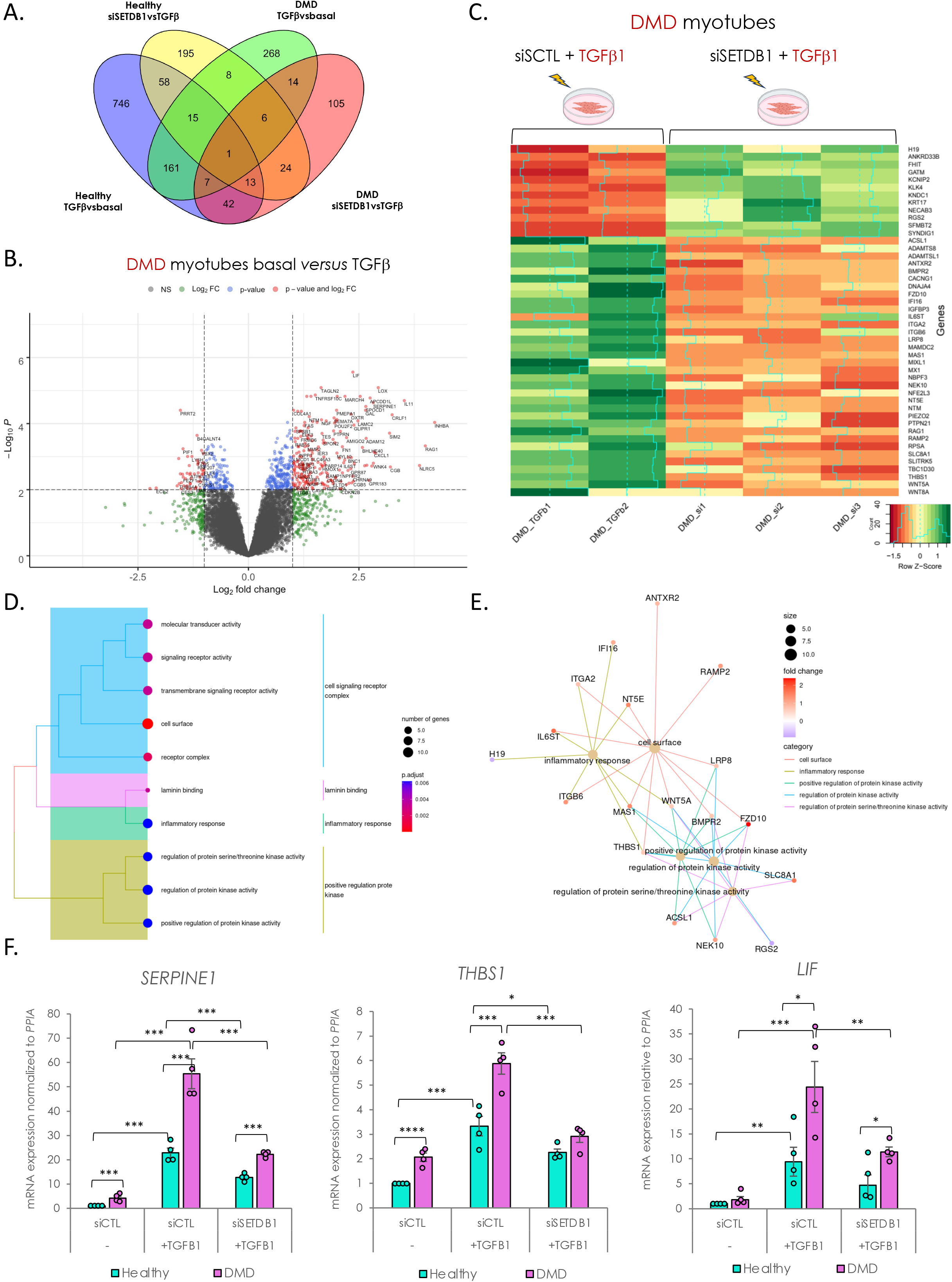
SETDB1 silencing in DMD myotubes leads to a decreased expression of TGFβ-dependent genes involved in ECM remodeling, receptor signaling transduction and inflammation. **A.** Venn diagram shows that a different set of TGFβ1-responsive genes were differentially expressed upon SETDB1 silencing in healthy and DMD myotubes. **B.** Volcano plot of differentially expressed genes in DMD myotubes +/- TGFβ1. **C.** Heatmap of expression level z-scores for DEGs in DMD myotubes for TGFβ *versus* TGFβ+siSETDB1 comparison. Some genes found to be increased upon TGFβ1 treatment decrease upon SETDB1 silencing (*IGFBP3, THBS1, IL6ST*). **D-E.** Enriched categories tree plot (D) and enriched gene-concept network for biological pathways deregulated for TGFβ *versus* TGFβ+siSETDB1 in DMD myotubes. **F.** mRNA levels of gene validated by RT-qPCR involved in ECM-remodeling (*THBS1, SERPINE1*) and inflammation (*LIF*). **For all panels:** Statistics were performed on ≥3 biological replicates and data are represented as average +/- SEM *p<0.05; **p<0.01; ***p<0.001 (unpaired Student’s t test).

Gene ontology (GO) analysis of the TFGβ-responsive genes in heathy myotubes showed a significant enrichment in terms related to the regulation of MAPK cascade, negative regulation of locomotion, cellular component movement, cell adhesion, SMAD protein phosphorylation, migration, motility, chemotaxis, vessel and nervous development (**Figure S4B**). While in DMD myotubes, GO showed that the main categories are related to cytokine-mediated signaling, extra cellular matrix (ECM) remodeling and organization, inflammation, SMAD protein phosphorylation and positive regulation of cell motility (**Figures 4B**, **S4C**), reminiscent of disease traits. Collectively, these data validated the responsiveness of both healthy and DMD myotubes to the TFGβ treatment.

Then, we have checked the DEGs in SETDB1 LOF condition, both in healthy and DMD myotubes treated with TGFβ. We found that SETDB1 LOF affected 320 genes in response to TFGβ in healthy myotubes (**Figure 4A**, top genes on **Table S3**). To determine the signature of the transcriptional changes, we applied gene set enrichment analysis (GSEA) and found that the SETDB1-dependent genes belong to ECM and cell surface categories (**Figure S4D**). Together, these data already suggest that SETDB1 LOF affects genes coding for secreted factors.

In DMD myotubes, SETDB1 LOF affected 212 genes in response to TFGβ (**Figure 4A, 4C**, top genes on **Table S4**), 120 were less abundant and 92 more abundant. In particular, SETDB1 LOF induced less abundant mRNAs associated with ECM remodeling, inflammation and fibrosis, including *THBS1* (*Thrombospondin-1*)*, Myostatin, BMPR2 (Bone Morphogenetic Protein Receptor type 2), ADAMTS8 (ADAM Metallopeptidase with Thrombospondin type 1 motif 8), MMP14 (Matrix Metallopeptidase 14), LIF (Leukemia Inhibitory Factor), SERPINE1 (Plasminogen Activator Inhibitor Type 1 or PAI1), WNT5A,* and *MAMDC2 (MAM domain containing 2)* (**Figures 4C**, **4F, S4E**). All these genes, gene categories and pathways are notably involved in the DMD disease traits. Furthermore, enrichment analysis for biological processes and gene network of biological pathway analyses highlighted that the SETDB1-dependent genetic programs in DMD myotubes in response to TFGβ are mainly involved in inflammation, signaling and cell surface (**Figure 4D-E**). SETDB1 KD alone, in absence of TGFβ, affects at a lesser extent the expression of the tested genes (**Figure S3D and S3H**).

Altogether, here we validated the efficacy of TGFβ to induce fibrotic and inflammation genetic programs and highlighted that SETDB1 silencing alters the transcription of many secreted factors (secretome) and ECM components, attenuating the deleterious effect of TGFβ overactivation in DMD myotubes.

### SETDB1 LOF in myotubes has a beneficial impact on the TGFβ−induced secretome: it improves muscle differentiation and reduces fibrosis

So far, our data showed that SETDB1 LOF in myotubes could potentially modulate the impact of TGFβ on surrounding cells and influence the process of muscle regeneration. To test this, we produced conditioned medium (CM) in healthy or DMD myotubes with or without SETDB1 LOF, treated or not with TGFβ (**Figure 5A**). Then, CM from healthy myotubes was applied on confluent healthy myoblasts, and CM from DMD myotubes on DMD myoblasts (**Figure 5A**).

**Figure 5:**
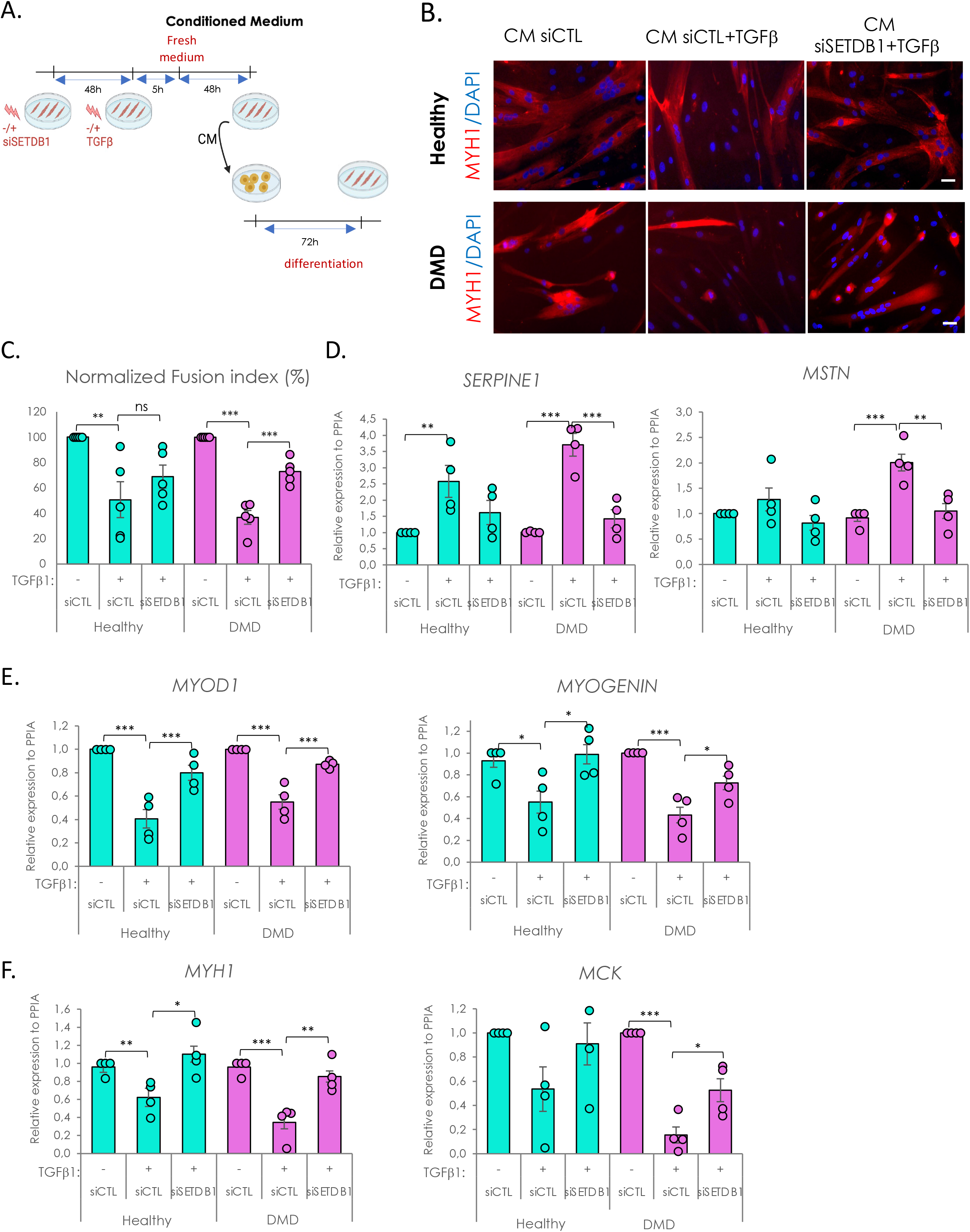
Secretome of SETDB1 deficient myotubes in response to TGFβ1 reduces the negative impact of TGFβ treatment on myoblast differentiation. **A.** Diagram of the conditioned medium experiments. **B.** Immunofluorescence of myosin heavy chain (red) and nuclei staining with DAPI (blue). Scale bar, 50 μM. **C.** Quantification of fusion index after 6 days of differentiation in the conditioned medium. **D.** RT-qPCR of fibrotic markers *SERPINE1* and *MSTN* after 3 days in the conditioned medium. **E.** RT-qPCR of early myogenic markers *MYOD1* and *MYOGENIN*. **F.** RT-qPCR of late myogenic markers *MYH1* and *MCK*. **For all panels:** Statistics were performed on ≥3 biological replicates (>100 nuclei for immunostaining quantification) and data are represented as average +/- SEM *p<0.05; **p<0.01; ***p<0.001 (unpaired Student’s t test).

As a control, we first checked whether CM could impact myoblast differentiation as compared to fresh differentiation medium and found no changes (**Figure S5A-C**). Thus, receiving myoblasts treated with fresh medium or CM supplemented with TGFβ1 showed a less efficient differentiation (**Figure S5A-C**). In absence of medium change, CM from TGFβ-treated myotubes completed impaired fusion, as expected, meaning that TGFβ1 is not consumed by the cells and remains high in the CM and though hides the myotube response (*18*).

We then included medium change after acute TGFβ1 treatment to study the intrinsic myotube pro-fibrotic response (**Figure 5A**). Our results showed that DMD myoblasts, but not healthy ones, have a decreased fusion index when differentiated with CM from TGFβ1-treated myotubes, but myoblasts incubated with CM from TGFβ1-treated and SETDB1 LOF myotubes show a more normal fusion index (**Figures 5B-C**). SETDB1 KD alone in myotubes, in absence of TGFβ, does not affect the fusion index of the myoblasts receiving the conditioned medium (**Figure S5E**).

The discrepancy between the effects of the CM in healthy versus DMD myoblasts could be due to the higher differentiation potential of the healthy myoblasts (**Figure S5D**).

Interestingly, the receiving myoblasts incubated with CM from TGFβ-treated and SETDB1 LOF myotubes showed a significant decrease in fibrotic markers *SERPINE1* and *MSTN* (*Myostatin*) (**Figure 5D**). Concomitantly, they displayed an increase in differentiation markers expression such as *MYOD1*, *Myogenin*, *MCK* (*Muscle Creatine Kinase*) and *MYH1* (**Figures 5E-F**). SETDB1 KD alone in myotubes, in absence of TGFβ, does not affect the expression of muscle differentiation genes in the myoblasts receiving the conditioned medium (**Figure S5F**).

Together, these data show that the TGFβ pathway activation could be attenuated by targeting SETDB1 in DMD patients. Thus, SETDB1 LOF in myofibers could be beneficial for the myofibers themselves but also to the differentiation of the surrounding myoblasts and potentially limit fibrosis.

## DISCUSSION

Tissue repair, such as muscle regeneration after injury, involves many cell types which communicate together through secreted molecules, the so-called secretome, to orchestrate replacement of damaged myofibers with new functional ones and re-establish tissue homeostasis. Thus, in addition to their contractile properties, myofibers have also a central role in cell-cell communication since they are able to secrete the so-called myokines especially during regeneration and differentiation, including dedicated vesicles acting on muscle adaptation to damage or exercise(*26*). This myofibers secretome plays important roles in intercellular communication of the muscle resident cells, i.e., MuSCs, myofibers themselves, the fibro-adipogenic progenitors (FAPs) and macrophages (*27*).

Although FAPs and macrophages have been described as major TGFβ sources, and as fibrosis and inflammation effectors, myofibers are also responsive to TFGβ1 and have been shown to participate in TGFβ/SMAD response during muscle regeneration *in vivo* (*28, 29*). TGFβ-induced fibrosis is a key pathological feature in muscle disease, such as DMD, and TGFβ overactivation worsens muscle degeneration by preventing proper repair. Moreover, even though myoblasts display a TGFβ/SMAD autocrine signal required for proliferation, TGFβ pathway prevents myoblast fusion by controlling actin remodeling and its activation decreases upon muscle differentiation. Interestingly, DMD muscle cellular models have been shown to recapitulate TGFβ/SMAD overactivation pathological feature *in vitro* associated with muscle defects and fibrosi*s* (*15, 16*). Hence, targeting TGFβ-induced pro-fibrotic response in DMD to slow down disease progression appears as a promising therapeutic approach.

TGFβ signaling nuclear endpoint mediated by the two major TGFβ downstream transcription factors SMAD2 and SMAD3, also involves many chromatin-modifying enzymes including SETDB1. SETDB1 regulates different cell fates including stemness and terminal differentiation (*2*). SETDB1 is also involved in fibrosis (*23*), inflammatory response and diseases (*5, 30–32*) and has been linked to TGFβ response in many non-muscle contexts (*19–23*). SETDB1 has been previously described by us and others as a major negative regulator of muscle terminal differentiation through repression of muscle gene expression in myoblast nuclei (*4, 33*). SETDB1 is responsive to Wnt signaling that is known to have a pro-differentiation effect on myoblasts and causes SETDB1 export to the cytoplasm allowing muscle gene de-repression. Interestingly, SETDB1 has also been shown to be a regulator of TGFβ/SMAD pathway in cancer (*19–22*) and pulmonary fibrosis (*23*) contexts. In this study, we have shown that SETDB1 nuclear localization follows TGFβ/SMAD activation in human muscle cells. SETDB1 is exported to the cytoplasm during muscle terminal differentiation, while TGFβ/SMAD pathway activation decreases, but it can be transiently relocalized in myotube nuclei in response to TGFβ pathway activation.

In the effort to identify new key players of TGFβ response whose dysregulation is a key driver of DMD progression, we hypothesized that SETDB1 could participate in TGFβ response in muscle and, though, could be involved in deregulation of TGFβ pathway in DMD. We found that SETDB1 is constitutively accumulated in DMD myotube nuclei correlating with phospho-SMAD3 high nuclear levels *in vivo* and *in vitro*. As expected, DMD myotubes are more responsive to TGFβ activation as compared to healthy ones. Interestingly, we were able to block SETDB1 nuclear localization by inhibiting intrinsic autocrine TGFβ signal in DMD myotubes. We also found that SETDB1 nuclear translocation is not dependent on SMAD3 protein and might involve phosphorylation events. We also performed a loss-of-function of SETDB1 and studied TGFβ-induced pro-fibrotic response. We found that healthy but especially DMD myotubes lacking SETDB1 display an attenuated response to TGFβ/SMAD pathway activation without changing phospho-SMAD3 levels. These results point to unprecedented link between SETDB1 and TGFβ response in muscle, and more especially in DMD context. Nonetheless, the exact mechanisms through which SETDB1 localization is controlled by either TGFβ or Wnt pathways are not yet understood.

To better understand how SETDB1 impacts TGFβ response in myotubes, we performed global transcriptomic analysis on healthy and DMD myotubes. We found that SETDB1 loss-of-function leads to a global decrease of RNA levels of gene coding for factors involved in ECM remodeling, inflammation and TGFβ and Wnt pathway signaling that respond to TGFβ. Since SETDB1 is known for its repressive action on gene expression, we made the hypothesis that SETDB1 could repress TGFβ inhibitors as already described in cancer context (*20*). Interestingly, we found that SETDB1 loss-of-function leads to an increased expression of the TGFβ inhibitor genes *SKIL* and *SMAD7*. The precise mechanism through which SETDB1 controls the expression of these genes remain unclear. Chromatin immunoprecipitation assays on SETDB1 and H3K9me3 marks would be required to confirm that SETDB1 represses TGFβ inhibitor expression but remain challenging in myotubes. In general, the mechanisms of action of SETDB1 at the chromatin level are well-documented (*34, 35*). Therefore, we can speculate that the enrichment of SETDB1 in the nucleus in DMD myotubes might affect chromatin accessibility and 3D genome organization, the points which would be interesting to address in future.

Finally, since most of SETDB1 direct or indirect targets are coding for secreted factors involved in fibrotic response and that TGFβ is known for its anti-differentiation role, we tested the effect of myotube secretome on myoblasts. Moreover, DMD muscle cells display fusion defects in some DMD models of hiPSC-derived myotubes and our model of immortalized myoblast-derived myotubes. Here, TGFβ-treated myotube secretome negatively impacts myoblast differentiation and more specifically fusion, but SETDB1 LOF in DMD myotubes has a beneficial cell non-autonomous effect on myoblast fusion. We show here the effect of SETDB1 on myotube secretome that affects environing cells. Targeting SETDB1 in post-mitotic myotubes does not seem to have deleterious effect on cell viability but its targeting in muscle tissue would be challenging since we do not know the impact on other cell types such as FAPs and macrophages. Nevertheless, we highlight interesting, secreted targets displaying aberrant expression in DMD myotubes that could play a role in deleterious environment of DMD. Moreover, some of deregulated targets have an unknown function such as *ANKRD33B* but could be involved in muscle regeneration process.

In summary, this study highlights the role of SETDB1 in post-mitotic muscle cells in adaptation to the environmental cues (**Figure S6**). Even though the mechanism(s) involved in SETDB1 regulation and the requirement of its enzymatic activity remains elusive, we unraveled a functional effect of its LOF on muscle cell differentiation and shed light on gene networks under SETDB1 control that would be worthy to study individually in the context of DMD. At last, we did not investigate Wnt and TGFβ interaction in this study but it seems that SETDB1 acts as a mediator of these pathway communication in muscle. Even though Wnt and TGFβ are described as having opposite effects on muscle terminal differentiation, some studies also described their collaboration in muscle (*36, 37*). Here, we showed *in vitro* the cell non-autonomous effect of SETDB1 inhibition in myotubes on the surrounding myoblast differentiation through the control of secreted factor expression induced by TGFβ/SMAD pathway. Nevertheless, it is unclear if this effect could be beneficial *in vivo* during regeneration since some of the targets have been described in literature for their double-edged effect on muscle regeneration and this could depend on the cell type involved. A fine-tuned balance has to be established in muscle tissue to allow a proper regeneration and SETDB1 could be involved in this mechanism by controlling muscle differentiation, but also by regulating myofiber response to environmental cues.

## MATERIALS & METHODS

### Establishment of stable cell lines and cell culture

The human induced pluripotent stem cell lines, control and DMD, generated from skin fibroblasts from Coriell (**Table 1**), were obtained from the Marseille Stem Cells platform in Marseille Medical Genetics laboratory and described in (*38*). IPSCs were maintained in culture in mTesR1 medium and dissociated using ReLeSRTM (StemcellTM). They were transfected with epB-Puro-TT-mMyoD and epB-Bsd-TT-FlagBaf60c2 by electroporation using the Neon Transfection System as described in (*15, 16*). Selection was performed using 5 μg/ml of Puromycin and 10 μg/ml of Blasticidin at the same time.

**Table 1:** Cell line list.

| *Cells* | *Reference* | *Patient* |
| --- | --- | --- |
| Induced pluripotent stem cells (IPSCs) | AG08C5 Coriell | Skin fibroblast, 1 year old healthy male |
|  | GM25313 | Skin fibroblast, 13-year-old DMD male patient, deletion ex45 |
| Immortalized myoblasts (hTERT/CDK4) (hMB) | AB1190C16PV | Paravertebral, 16-year-old healthy male |
|  | AB1167C20TFL | Fascia lata, 20-year-old healthy male |
|  | AB1023DMD11Q | Quadriceps, 11-year-old DMD male |
|  | AB1071DMD13PV | Paravertebral, 13-year-old DMD male |

Human control and DMD immortalized myoblasts were obtained from AFM-MyoLine. Immortalization was performed as described in: (*24*), using human telomerase-expressing and cyclin-dependent kinase 4-expressing vectors. They were cultured on Gelatin-coated plates and propagated in DMEM high glucose GlutaMAX™ (Invitrogen, 61965-026)/Medium 199 (Invitrogen, 41150020) 4:1 mixture, supplemented with 20% FBS (Sigma, F7524), 50 μg/mL Fetuin, 5 μg/mL insulin, 0.5 ng/mL bFGF, 5 ng/mL hEGF and 0.2 μg/mL Dexamethasone (Sigma D4902).

### Myogenic differentiation of iPSCs and human immortalized myoblasts

iPSCs stably expressing MyoD and Baf60C Tet-ON inducible transgenes were propagated in mTeSR1 on Matrigel-coated wells. To induce myogenic differentiation starting from iPSC colonies, doxycycline (200 ng/mL) was added in cells maintained in mTeSR1 (day 0). After 24 h of treatment with doxycycline (day1), cells were dissociated as single cells using TrypLE and plated 25.10^3^/cm2 in growth media (GM) (knockout DMEM (Invitrogen) supplemented with 1 mM L-glutamine, 20% knockout serum replacement medium (KOSR, Invitrogen), Glutamax 1X, 0.1 mM nonessential aminoacids (NEAA, Invitrogen), 50 U/mL streptomycin (Invitrogen) plus hES cell recovery supplement (10 μM) (Stemgent) and doxycycline. On day 3, GM medium was switched to differentiation media (DM) (knockout serum-free DMEM containing 1X Insulin-Transferrin-Selenium (ITS) (SIGMA)) supplemented with doxycycline until day 7.

Human control and DMD immortalized myoblasts were grown at > 80% confluence and the medium was switched for DM media.

### Myotube transfection with siRNAs

Three days-differentiated myotubes cultured in differentiation medium (DM) were transfected with siRNAs at a final concentration of 70 nM, employing the Lipofectamin™ RNAiMAX transfection agent (Invitrogen™, #13778100). Cells were kept 2 days in transfection medium before switching with fresh DM. We used the ON-TARGETplus Human SETDB1 siRNA (DharmaconTM; L-020070-00-0010); *SMAD3* siRNA (Sens: GUGUGAGUUCGCCUUCAAUAU; Antisens:

AUAUUGAAGGCGAACUCACAC), and the ON-TARGETplus Non-targeting Control Pool (DharmaconTM; D-001810-10) as a scrambled control siRNA.

### Immunofluorescence on cells and tissue

Muscle histological sections were obtained from muscle biopsies of 15-years old DMD patient paravertebral muscle and a healthy 17-years old individual. Frozen samples were fixed in ice-cold acetone for 1 min and then air-dried. They were blocked in 4% BSA-PBS solution for 45 min at room temperature (RT) and then incubated with the primary antibodies at 4°C overnight. Secondary antibodies were applied for 45 min at RT in the dark before mounting.

Cells were grown on Matrigel- and Gelatin-coated coverslips for iPSCs and human myoblasts respectively. They were fixed with 4% PFA for 20 min and they were saturated with 50 mM NH4Cl for 10 min at RT. Then, they were permeabilized with 0,5% Triton X-100 for 5 min. Cells were washed 3 times with PBS between each steps. Cells were next incubated with primary antibodies at the concentration indicated in the antibody list below. Alexa-488 or 555 or 647 were used as secondary antibodies (Invitrogen). Nuclei were counterstained with DAPI.

Images were acquired with a Leica DMI-6000B microscope and analysis was performed using Fiji software. In-house macro was used to performed nuclear and cytoplasmic signal quantification. Acquisition of iPSCs-derived myotubes was performed using ImageXpress Micro High Content Screening System. All the conditions were acquired with identical settings and were analyzed with an in-house macro on ImageJ/Fiji to quantify nuclear and cytoplasmic signal. The used antibodies are listed on **Table 2**.

**Table 2:** Antibodies list.

| *Antibody* | *reference* | *Use & dilution* |
| --- | --- | --- |
| SETDB1 (rabbit) | Santa Cruz sc-66884 | Immunofluorescence (1:200) |
| SETDB1 (mouse) | Abcam ab107225 | Western blot (1:1000) |
| SETDB1 (mouse) | Thermo Fisher Scientific MA5-15722 | Immunofluorescence (1:200) |
| Myosin heavy chain, MYH1E | DSHB | Immunofluorescence (1:10), Western blot (1:200) |
| Phospho-SMAD3 (rabbit) | Abcam ab52903 | Immunofluorescence (1:200), Western blot (1:1000) |
| SMAD3 (mouse) | Proteintech 66516-1-Ig | Western blot (1:1000) |
| PAI1 (rabbit) | Proteintech 13801-1-AP | Western blot (1:1000) |
| RNA polymerase II (rabbit) | Abcam ab5095 | Western blot (1:1000) |
| Lamin A/C (rabbit) | Proteintech 10298-1-AP | Western blot (1:1000) |
| H3 (C-16) (goat) | Santa Cruz sc-8654 | Western blot (1:1000) |

### RNA and Quantitative Reverse Transcription-PCR (RT-qPCR)

Total RNA was extracted using RNeasy micro-kit (Qiagen) following manufacturer’s procedures. DNase (Qiagen) treatment was performed to remove residual DNA. 1 μg of total RNA was reverse transcribed with High-Capacity cDNA Reverse Transcription Kit (Applied Biosystems). Real-time quantitative PCR was performed to analyze relative gene expression levels using SYBR Green Master mix (Applied Biosystems) following manufacturer indications. Relative expression values were normalized to the housekeeping genes mRNA *PPIA* or *UBC*. Primers are listed in **Table 3**.

**Table 3:**
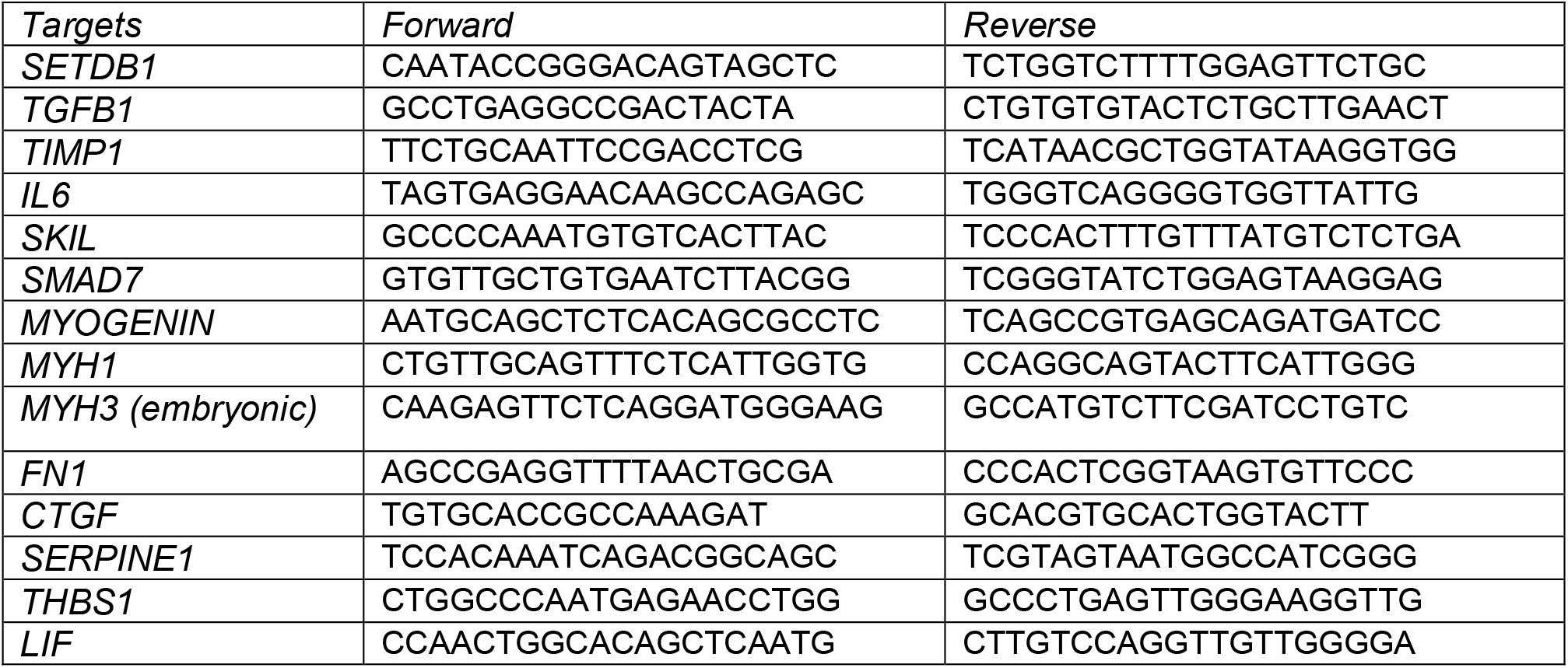
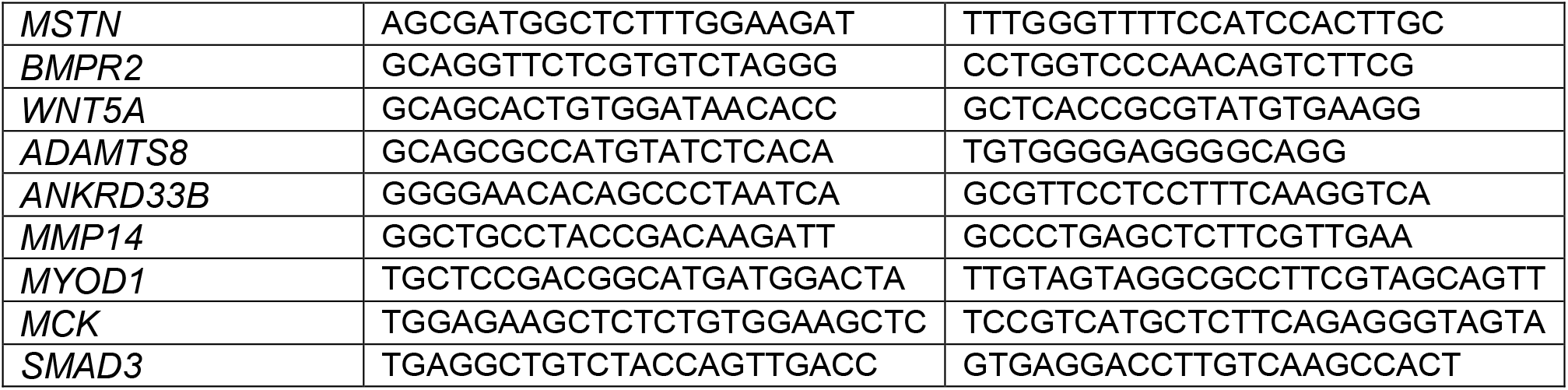
Primer list.

### Nuclear and cytoplasmic fractionation

Cells were scraped directly in 3 volumes of buffer A (20 mM HEPES pH7.0, 0.15 mM EDTA, 0.15 mM EGTA, 10 mM KCl) supplemented with 0.15 mM Spermine, 0.5 mM Spermidine, protease inhibitor cocktail 1X (SIGMAFAST^TM^) and phosphatase inhibitor cocktail 1X (Sigma). Cells were lysed with 0.5% NP-40 and mixed gently by inversion. 0.88 volume of buffer SR (50 mM HEPES pH7, 0.25 mM EDTA, 10 mM KCl, 7% sucrose) supplemented with Spermine, Spermidine, protease and phosphatase inhibitor cocktails was added before centrifugating the cells for 5 min at 2000 x g at 4°C and the supernatant was collected as the cytoplasmic fraction. The core nuclei pelleted at 2000 x g was resuspended in 1 volume of Low Salt buffer (20 mM Tris pH7.65, 25% glycerol, 1.5 mM MgCl2, 0.2 mM EDTA, 20 mM NaCl) with 3X of protease and phosphatase inhibitor cocktails. One volume of High Salt buffer (same as Low Salt but with 900 mM NaCl) supplemented with 5 mM ATP while vortexing. The nuclei samples were kept on ice for 30 min and mix by inversion every 5 min. 1 volume of sucrose buffer (20 mM Tris pH7.65, 60 mM NaCl, 15 mM KCl and 0.34 M sucrose) was added before treating the cells with 0.0025 U/μL of MNase for 10 min at 37°C and 1 mM CaCl2. Reaction was stopped with 4 mM EDTA. Samples were next sonicated for 10 min and ultracentrifugated at 40000 rpm at 4°C for 30 min. Supernatant was collected as nuclear fraction.

### Western blot

Cells were lysed in RIPA buffer (20 mM Tris pH 7.65, 150 mM NaCl, 0.1% SDS, 0.25% NaDoc and 1% NP-40) supplemented with protease inhibitor 1X (SIGMAFAST^TM^) and phosphatase inhibitor 1X (Sigma) and kept on ice for 30 min. Cell lysates were sonicated at 4°C for 10 min (30 sec ON, 30 sec OFF) at medium frequency (Bioruptor Diagenode). Then, the lysates were centrifugated for 10 min at 4°C at maximum speed and the supernatants were kept as the samples. Extracts were resolved on pre-cast polyacrylamide gel cassettes (NuPAGE® 4-12% Bis-Tris) (Invitrogen) and 1X NuPAGE MOPS SDS Running Buffer and transferred into nitrocellulose membrane (Amersham) in 20 mM phosphate transfer buffer (pH 6.7). Membrane was blocked in 5% skim milk in PBST Buffer (1X PBS, 0.2% Tween 20) and incubated overnight at 4°C with the primary antibody (see Table of antibodies). Membranes were washed twice 5 minutes in PBST, incubated with appropriate secondary antibody IRDye® (Li-Cor) in PBST, washed twice 10 minutes in PBST, once 10 minutes in PBS and then imaged on Odyssey® Imaging System. The used antibodies are listed on **Table 2**.

### Transcriptome analysis and bioinformatics

RNA was isolated as described above. Two or three independent biological replicates were sequenced depending on the cell conditions. Libraries were generated using the Ion AmpliSeq^TM^ Transcriptome Human Gene Expression Kit (A24325; Ion Torrent^TM^). Sequencing was performed on an Ion S5 sequencer (Ion Torrent^TM^). The reads were analyzed using the plugin AmpliseqRNA of Torrent Suite (release 5.10) and mapped on the panel of human AmpliseqRNA which reveals the expression of 20813 genes. Out of these 18185 genes were uniquely mapped on human reference gene names (retrieved from ENSEMBL_102 release).

Using the raw count Table generated, genes having low counts were filtered out based on the log2 of Counts Per Million (logCPM). We kept genes that have at least a 1 logCPM mean expression level in at least one of the experimental conditions, leaving 12 060 expressed genes for all the downstream analyses. The differential expression analysis of the filtered data was performed using the edgeR package of R and the limma-trend method described in (*39*). For each experimental setting Differentially Expressed Genes (DEGs) were further analyzed for all the available enrichments. Enrichment analysis and visualization of the DEGs were performed using the clusterProfiler R package (*40*), using Gene Ontologies, DOSE, the MSigDB database and GSEA (*41*).

### Statistical analyses

Statistical analyses were carried out using Excel and R. Data are represented as mean +/- SEM as described in the figure legend. Graphs are prepared using Excel and R. Double tail *t* test was used for statistical analysis, *\*p<0.05, **p<0.01, ***p<0.001*.

## Acknowledgements

We thank members of the Ait-Si-Ali lab and Epigenetics and Cell Fate department for helpful discussions during the group and department meetings and critical reading of the manuscript. We thank Dr Vincent Mouly from the Institute of Myology and MyoLine platform, as well as Dr Frédérique Magdinier, Marseille Stem Cells platform in Marseille Medical Genetics laboratory, for generous sharing of biological material. We warmly thank Dr Capucine Trollet for her advice, generous sharing of biological material, technical and critical help. We warmly thank Anna Moles for NGS data thanks to the “Project CTN01_00177_888744 for the creation of a multiregional infrastructure (Italian regenerative medicine infrastructure IRMI) for the development of advanced therapies aimed at organ and tissue regeneration”. We thank the (EPI)2 Imaging platform and the EpHISTain platform - UMR7216 Epigenetic and Cell Fate Centre, for access to instruments and technical advice.

## Author Contributions

Conceptualization, S.A.S, F..L.G and S.A; Methodology A.G, M.Z, R.P, Ma.M, E.B, L.D.M, C.B, V.J, S.A., My.M, E.N, A.B and S.A.S. Software, C.B, S.P; Formal Analysis, C.B., S.P and A.G; Investigation, A.G, M.Z, L.D.M, C.B, V.J, F.L.G, S.A and S.A.S; Data Curation, C.B.; Writing – Original draft, A.G and S.A.S; Writing – Review & Editing, A.G, M.Z, L.D.M, C.B, V.J, E.N, S.A and S.A.S; Supervision, S.A and S.A.S; Funding Acquisition, S.A.S, F.L.G; Project Administration, S.A.S.

## Data and Code Availability

All data needed to evaluate the conclusions in the paper are present in the paper and/or the Supplementary Materials. Data corresponding to all the experiments described in this study are deposited as raw BAM files at the European Nucleotide Archive (ENA) under the accession number PRJEB63300.

## Supplementary Materials

**Figure S1:**
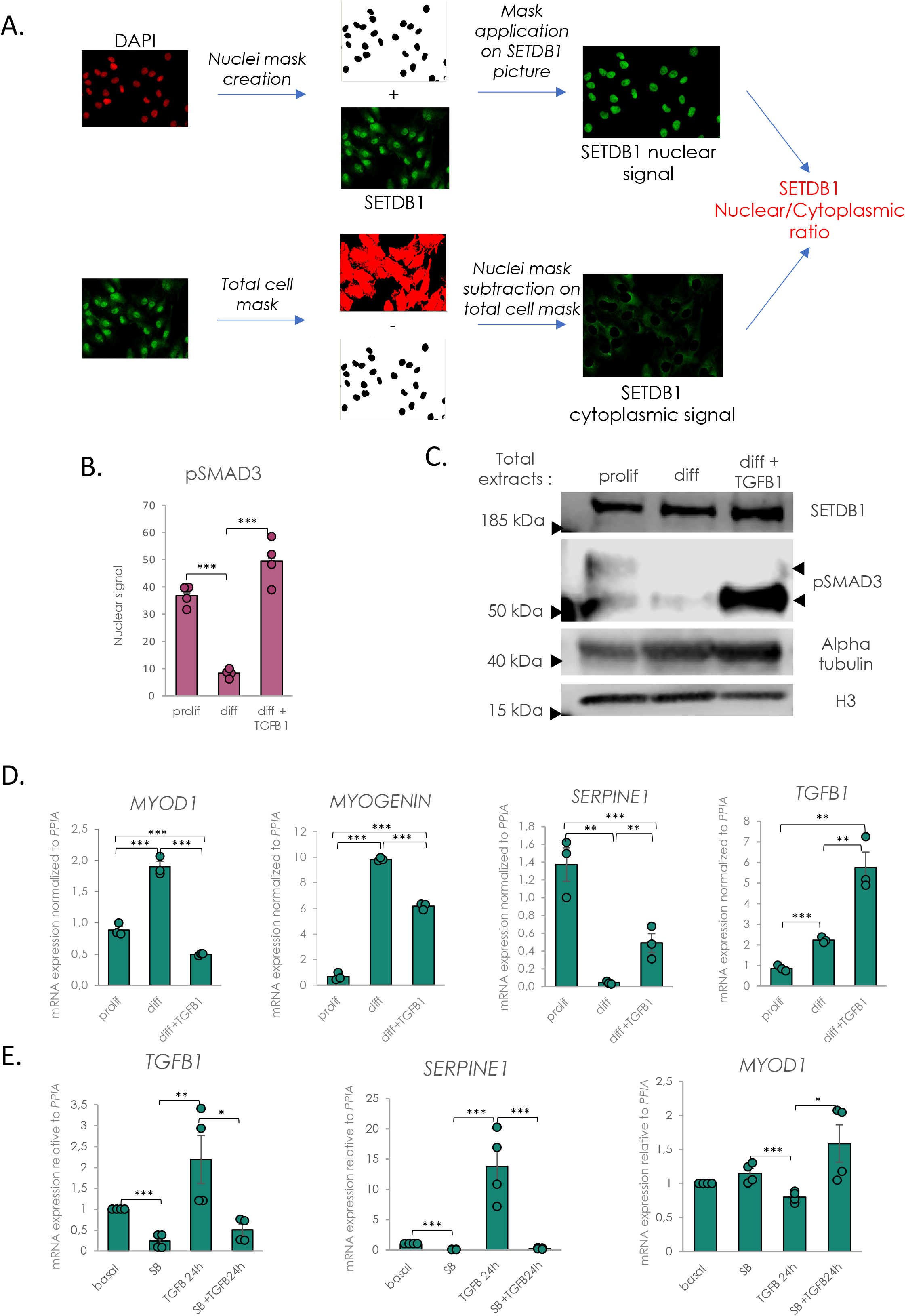
SETDB1 localization depends on TGFβ/SMAD pathway activation during muscle terminal differentiation. **A.** Scheme of microscopy analysis and nuclear/cytoplasmic signal quantification. Nuclear masks were setup from DAPI pictures and applied on SETDB1 pictures and total cell masks were selected on SETDB1 pictures. Cytoplasmic quantification was performed by subtracting nuclei mask from total cell masks. **B.** Quantification of phospho-SMAD3 nuclear fluorescence in proliferating myoblasts, myotubes treated or not with TGFβ1. **C.** Protein levels of SETDB1 and phospho-SMAD3 in total extracts of myoblasts and myotubes treated or not with TGFβ1. **B.** RT-qPCR of early myogenic (*MYOD1, Myogenin*) and TGFβ-related genes (*TGFB1, SERPINE1*) in myoblasts and myotubes treated or not with TGFβ1. **E.** RT-qPCR of early myogenic marker *MYOD1* and TGFβ-related genes (*SERPINE1, TGFB1*) in myotubes treated or not with TGFβ1 and/or its inhibitor SB-431542. **For all panels:** Statistics were performed on ≥3 biological replicates (>100 nuclei for immunostaining quantification) and data are represented as average +/- SEM *p<0.05; **p<0.01; ***p<0.001 (unpaired Student’s t test).

**Figure S2:**
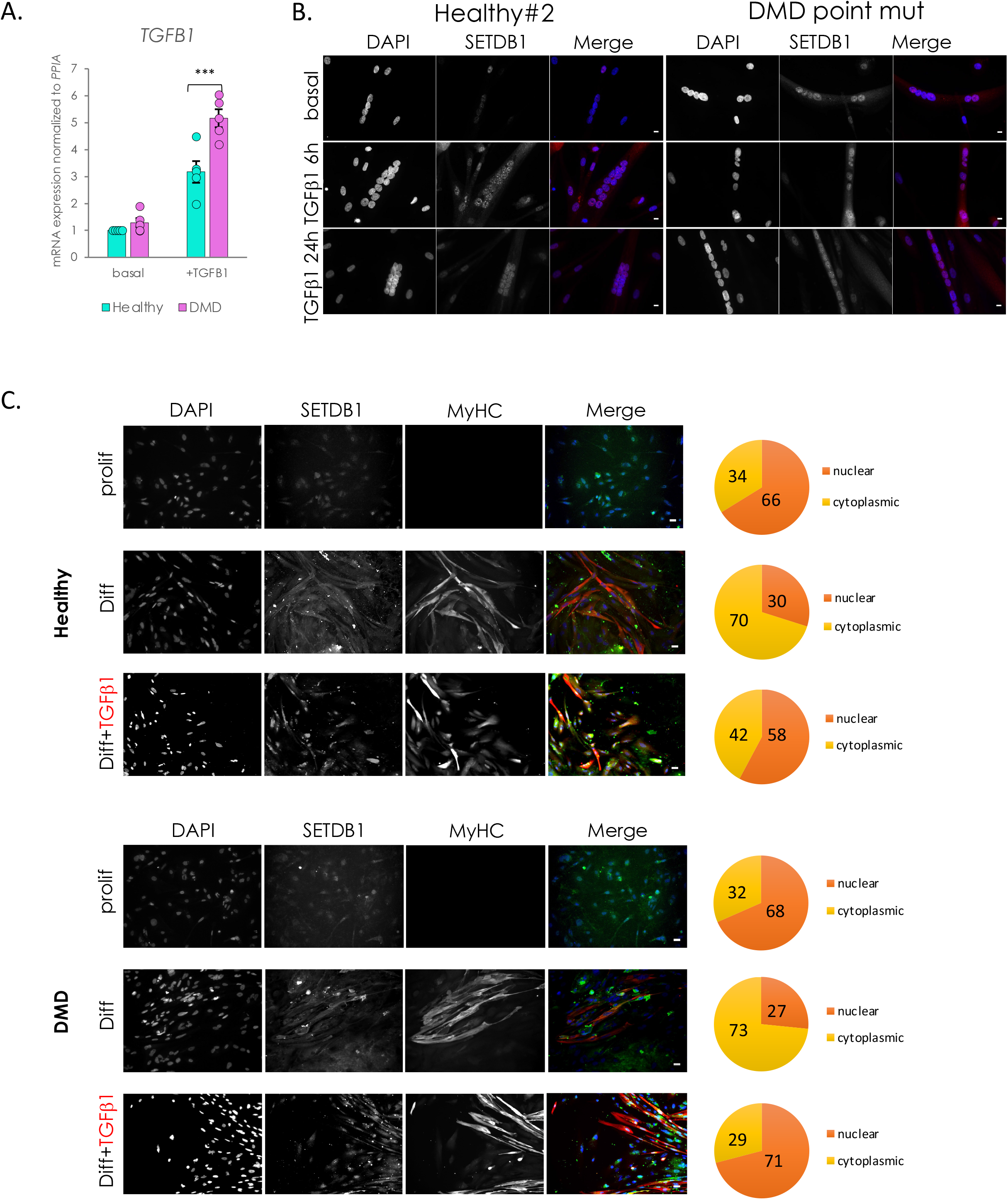
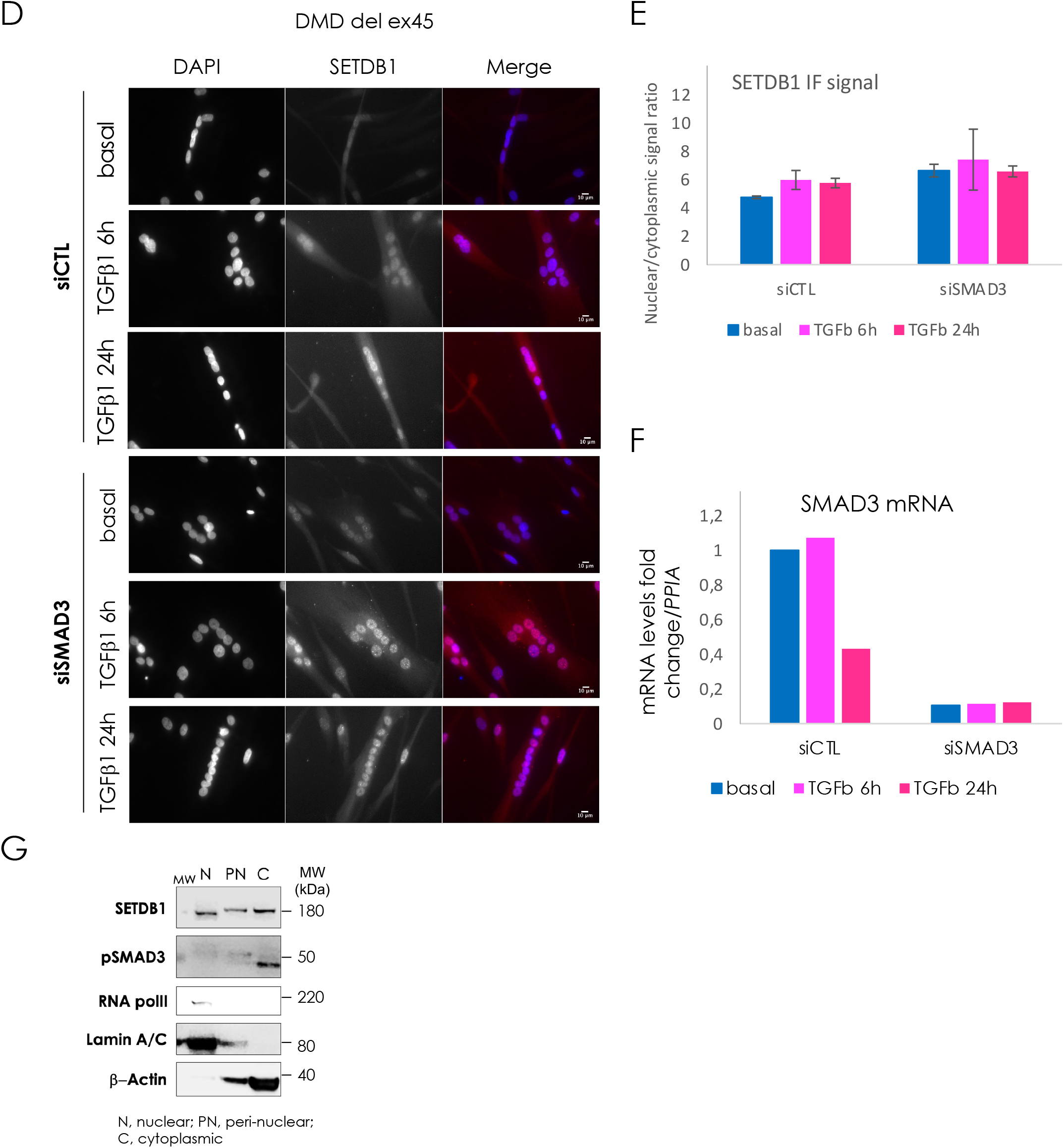
SETDB1 translocate into muscle cell nuclei in response to TGFβ/SMAD pathway activation and show more persistent nuclear signal in DMD myotubes irrespectively of the type of DMD mutation. **A.** RT-qPCR of *TGFB1* shows a higher response of DMD myotubes to TGFβ1 treatment as compared to WT cells. **B.** Immunostaining of SETDB1 (red) in healthy#2 and DMD point mutation myotubes. Nuclei were stained with DAPI (blue). Scale bar, 10 μM. **C.** SETDB1 and MyHC immunostaining in proliferating muscle cells and in myotubes treated or not with TGFβ1 derived from iPSCs of healthy or DMD individuals. Nuclei were stained with DAPI (blue). Scale bar, 10 μM. Diagrams represent the percentage of SETDB1 signal intensity measure in nuclei and cytoplasm of the cells. **D-F.** DMD del ex45 differentiating myotubes (48 h differentiation) were transfected with control scrambled (siCTL) or SMAD3 siRNA (siSMAD3). 2 days later, myotubes were treated with TGFβ and after 1 day, myotubes were subjected to different assays, as follows: **F.** Immunostaining of SETDB1 (red), and nuclei staining with DAPI (blue). Scale bar, 10 μm. **G.** Quantification of SETDB1 (n=2) nuclear/cytoplasmic IF signal ratio. **H.** RT-qPCR of *SMAD3* mRNA in control siRNA (siCTL) and siRNA against SMAD3 (siSMAD3) conditions to check the siRNA efficiency (representative of n=2). **G.** Western blot showing SETDB1 and pSMAD3 protein migration profiles in nuclear (N), peri-nuclear (PN) and cytoplasmic (C) fractions of healthy myotubes. RNA polymerase II and Lamin A/C were used as controls for nuclear fraction and beta-Actin as a control of cytoplasmic fraction. **For all panels:** Statistics were performed on ≥3 biological replicates (>100 nuclei for immunostaining quantification) and data are represented as average +/- SEM *p<0.05; **p<0.01; ***p<0.001 (unpaired Student’s t test).

**Figure S3:**
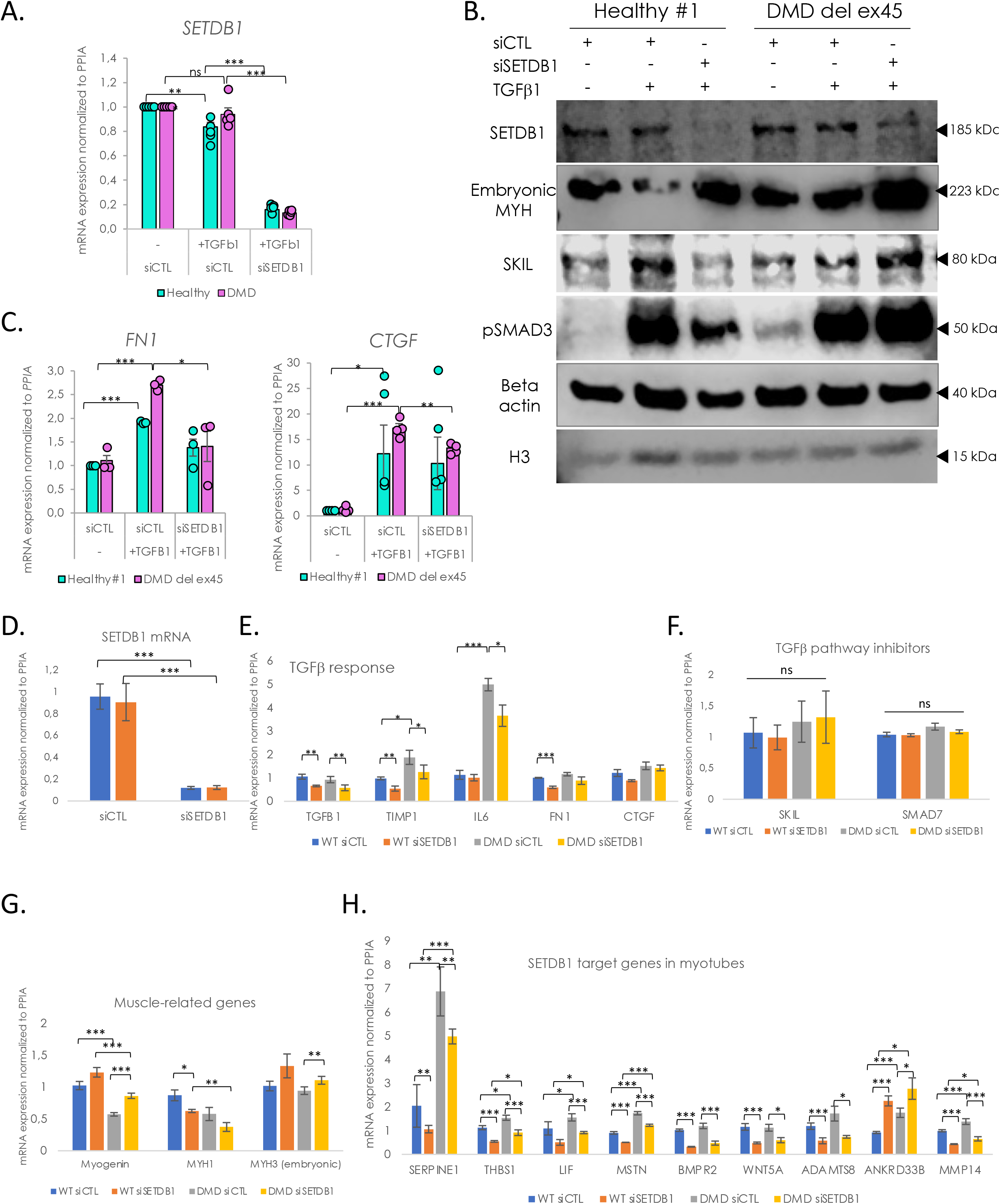
Efficient siRNA-mediated SETDB1 knockdown in TGFβ-treated DMD myotubes leads to a decrease in TGFβ target gene expression and an increase in pro-myogenic factors *SKIL* and *MYH3*. **A.** RT-qPCR of *SETDB1* shows an acute decrease (>80%) of *SETDB1* expression upon siRNAs-mediated silencing. *SETDB1* expression decreases upon TGFβ1 treatment in healthy myotubes but not in DMD myotubes. **B.** Western blot showing protein levels of SETDB1, SKIL, embryonic MYH and phospho-SMAD3 in healthy *versus* DMD myotubes upon SETDB1 silencing and TGFβ1 treatment. **C.** RT-qPCR of TGFβ/SMADs pathway known targets *FN1* and *CTGF* in healthy and DMD myotubes +/- siSETDB1 +/- TGFβ1. TGFβ-related gene expression is decreased in SETDB1-silenced *versus* siCTL DMD myotubes in response to TGFβ1. **D-H.** RT-qPCR of *SETDB1* (**D**), TGFβ known target genes *TGFB1*, *TIMP1*, *IL6*, *FN1* and *CTGF* (**E**), of TGFβ pathway inhibitors *SKIL* and *SMAD7* (**F**), muscle differentiation genes *Myogenin*, *MYH1* and *MYH3* (**G**) and of SETDB1 target genes such as *SERPINE1*, *THBS1*, *LIF*, *MSTN*, *BMPR2*, *WNT5A*, *ADAMTS8*, *ANKRD33B* and *MMP14* (**H**) in healthy and DMD myotubes +/- siSETDB1 in basal condition (without TGFβ treatment). **For all panels:** Statistics were performed on ≥3 biological replicates and data are represented as average +/- SEM *p<0.05; **p<0.01; ***p<0.001 (unpaired Student’s t test).

**Figure S4:**
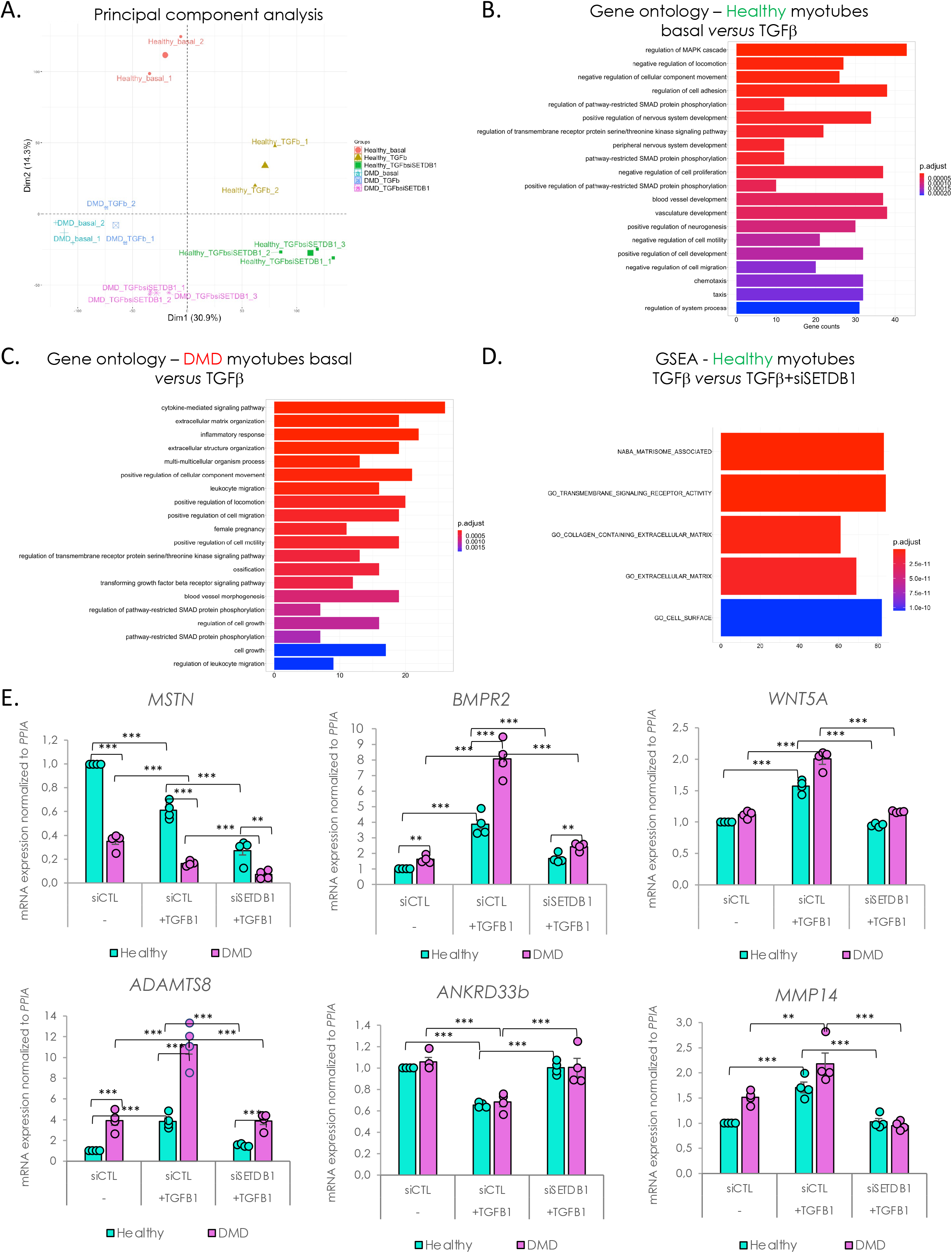
Healthy and DMD myotubes respond differently to TGFβ/SMAD pathway activation but display some SETDB1 target gene signatures in common. **A.** Principal component analysis of the filtered (see Materials & Methods) RNA-seq data, depicting a well grouping of all the samples for each of the 6 experimental conditions of the study. The variability captured by the first PC corresponds to the healthy vs. DMD conditions and the one of the second PC to the TGFβ treatment. **B.** Gene Ontology enrichment of DEGs from healthy myotubes basal *versus* TGFβ comparison. **C.** Gene Ontology enrichment of DEGs from DMD myotubes basal *versus* TGFβ comparison. **D.** Gene Set Enrichment Analysis (GSEA) of DEGs from healthy myotubes TGFβ +/- siSETDB1. **E.** Validation by RT-qPCR of genes coding for proteins involved in ECM-remodeling (*ADAMTS8, MMP14*), TGFβ/BMP and Wnt pathway (*MSTN, BMPR2, WNT5A*) and unknown function but predicted as regulator of muscle differentiation (*ANKRD33B*). **For all panels:** Statistics were performed on ≥3 biological replicates and data are represented as average +/- SEM *p<0.05; **p<0.01; ***p<0.001 (unpaired Student’s t test).

**Figure S5:**
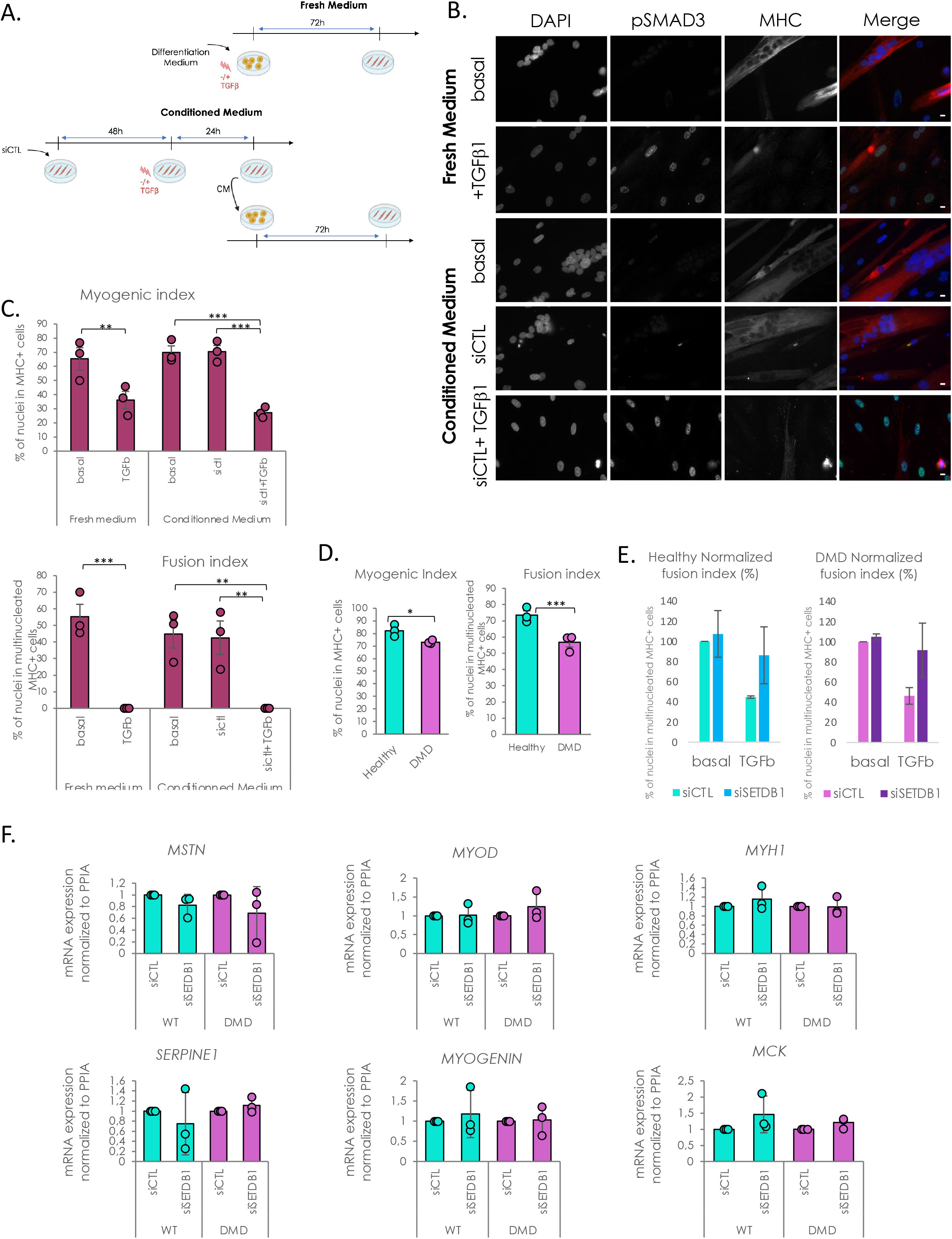
TGFβ/SMAD pathway activation leads to fusion defects in muscle cells. **A.** Diagram of the experimental design. **B.** Immunofluorescence of pSMAD3 (green) and MHC (red) in myoblasts differentiated in fresh or conditioned medium +/- TGFβ1. Nuclei were stained with DAPI (blue). Scale bar, 10 μM. **C.** Myogenic and fusion index of the myoblasts differentiated in fresh or conditioned medium +/- TGFβ1. **D.** Raw myogenic and fusion index in basal conditioned medium show difference in differentiation rate between healthy and DMD myotubes. B. Quantification of fusion index after 6 days of differentiation in conditioned medium produced by healthy or DMD myotubes +/- siSETDB1 +/- TGFβ1. **F.** RT-qPCR of pro-fibrotic genes, *MSTN* and *SERPINE1*, and muscle-related genes *MYOD1*, *Myogenin*, *MYH1* and *MCK* after 3 days of differentiation in conditioned medium produced by healthy or DMD myotubes +/- siSETDB1. **For all panels:** Statistics were performed on ≥3 biological replicates (>100 nuclei for immunostaining quantification) and data are represented as average +/- SEM *p<0.05; **p<0.01; ***p<0.001 (unpaired Student’s t test).

**Figure S6:**
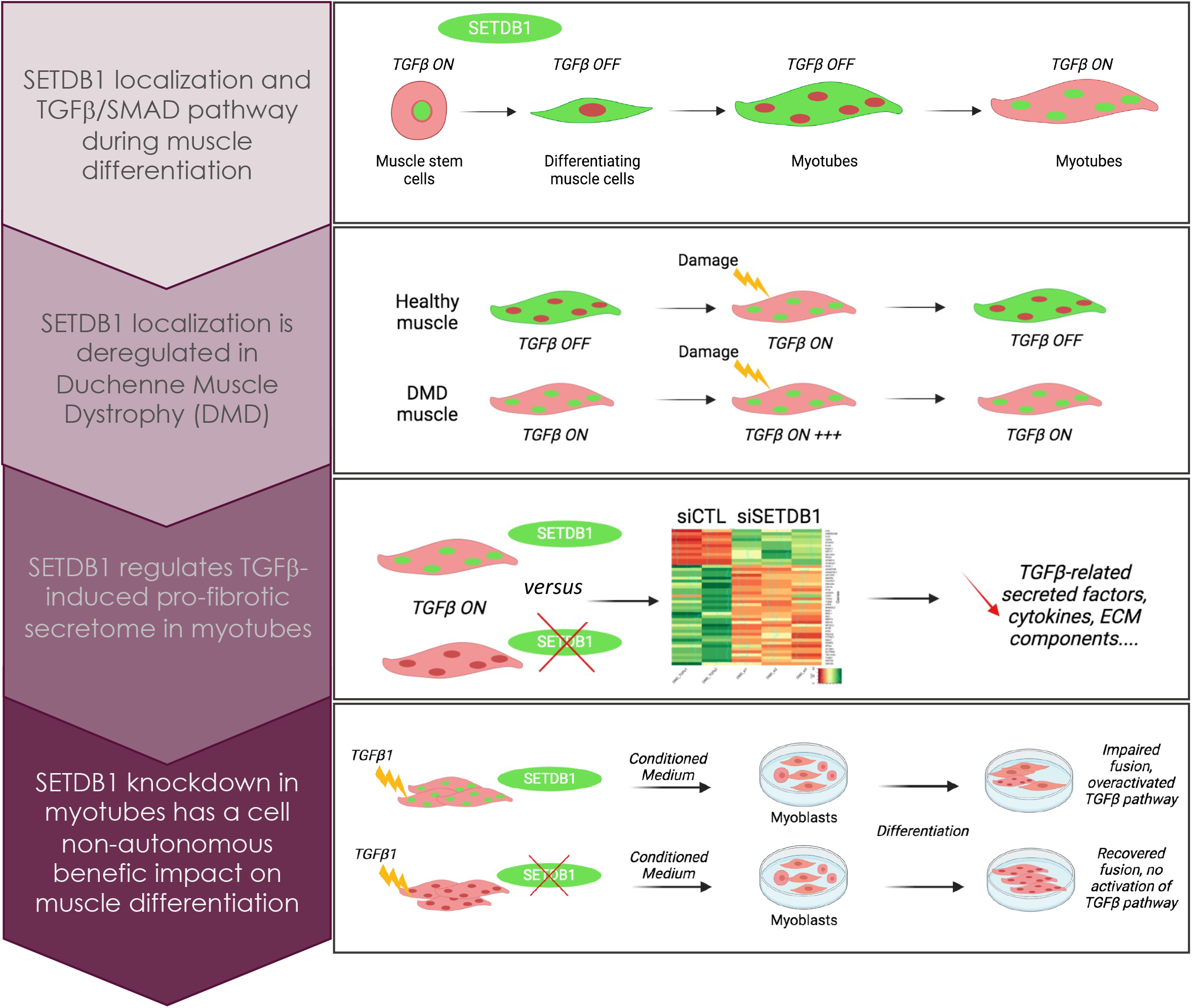
Graphical Abstract. - TGFβ induces nuclear accumulation of SETDB1 in healthy myotubes
- SETDB1 is enriched in DMD myotube nuclei with intrinsic TGFβ pathway overactivation
- SETDB1 LOF in DMD myotubes attenuates TGFβ-induced pro-fibrotic response
- Secretome of TGFβ-treated DMD myotubes with SETDB1 LOF is less deleterious on myoblast differentiation

**Table 1:** Top DEGs + *versus* – TGFB1 in healthy myotubes.

| Genes | LogFC | Adj p-value | Genes | LogFC | Adj p-value | Genes | LogFC | Adj p-value |
| --- | --- | --- | --- | --- | --- | --- | --- | --- |
| TAGLN2 | 2,117414122 | 0,0000029 | FZD1 | -1,709895607 | 0,000820576 | CYGB | 1,91232727 | 0,004636356 |
| PMAIP1 | 1,822825158 | 0,0000295 | CPT1A | 1,892601801 | 0,000847677 | CHRNA9 | 2,131251441 | 0,004655553 |
| SPSB1 | 1,954444533 | 0,0000431 | FLRT3 | -1,666029718 | 0,000907678 | HES6 | 1,511256537 | 0,004686457 |
| LEFTY2 | 2,378015571 | 0,0000633 | SPOCD1 | 2,072190972 | 0,000955143 | BHLHE40 | 1,577516443 | 0,004688974 |
| MYL10 | 2,775733677 | 0,000102766 | KY | 1,741592873 | 0,000987349 | DUSP15 | 1,504729237 | 0,004688974 |
| NGF | 1,657626522 | 0,000102766 | MMP24 | 1,785794877 | 0,001060517 | PRTG | -1,939105695 | 0,004688974 |
| HSD11B2 | 2,056381007 | 0,00014734 | PTPRN | 1,572910987 | 0,001060517 | TAS2R50 | -1,617874683 | 0,004707551 |
| CPEB1 | 2,008260893 | 0,00014734 | SPHK1 | 1,553357986 | 0,001060517 | CAPG | 1,617433777 | 0,004761555 |
| GRIK3 | 1,897772958 | 0,00014734 | SLC6A15 | -1,604281321 | 0,001060517 | IL15 | 1,632420492 | 0,004800381 |
| CAMK2B | 1,66141848 | 0,00014734 | RGS10 | 1,775216817 | 0,001091293 | RCSD1 | 1,50536248 | 0,005066626 |
| LIF | 1,609459933 | 0,00014734 | ACAP1 | -1,518147984 | 0,001091293 | BCHE | -1,514123565 | 0,00508091 |
| FAS | 1,65594723 | 0,000176389 | CDH22 | 1,952879439 | 0,001160065 | HAPLN1 | -1,939131798 | 0,005275934 |
| COMP | 3,079879046 | 0,000193172 | VDR | 2,38037933 | 0,001185628 | TRAPPC11 | -1,683727235 | 0,005288437 |
| KRT17 | 2,249228221 | 0,000193172 | MADCAM1 | 2,012788312 | 0,001218815 | E2F2 | -1,769263425 | 0,005304981 |
| CYSLTR1 | -1,686154285 | 0,000193172 | IL11 | 1,720981756 | 0,00125013 | ABHD1 | -1,604574292 | 0,005424992 |
| EFNA5 | -1,539212907 | 0,000193172 | ATF3 | 1,505914322 | 0,001406572 | EDN3 | -2,42948784 | 0,005452562 |
| CBFA2T3 | 1,585761128 | 0,00021232 | BNC1 | 1,727769999 | 0,001463008 | KCNG3 | -2,006427044 | 0,005469823 |
| TINAGL1 | 2,040560934 | 0,000214261 | UGT8 | -2,091670121 | 0,001463008 | MT2A | 2,078060952 | 0,005630917 |
| ST6GALNAC2 | 2,621131534 | 0,000278225 | MDGA1 | -1,57495199 | 0,001500196 | MMRN1 | -5,596523239 | 0,005630917 |
| MYL7 | 1,928998074 | 0,000278225 | LRRN3 | -2,023789769 | 0,001564602 | NPPB | 2,22275891 | 0,005782479 |
| RAB11FIP4 | 1,68799875 | 0,000278225 | APCDD1L | 1,573817321 | 0,001603986 | SFRP1 | -1,766514568 | 0,005851596 |
| CADM2 | -1,756947008 | 0,000278225 | LEFTY1 | 2,948081363 | 0,001646694 | NUPR1 | 2,251896144 | 0,006190195 |
| KCNH3 | 1,526606981 | 0,000294724 | INSC | 1,984068026 | 0,001654329 | MAPK10 | -1,765559007 | 0,006190195 |
| SH2D4A | 1,542140036 | 0,000304292 | FGFBP3 | -1,960631592 | 0,001750743 | TENM1 | -1,823947329 | 0,006245981 |
| BMP8A | 1,66627264 | 0,000311971 | LRP2 | -2,613936968 | 0,001789815 | ABCA7 | -1,6412153 | 0,006245981 |
| UBASH3B | 1,556460437 | 0,000311971 | KCNK3 | 2,416533797 | 0,001890051 | ADAMTS3 | -1,554488887 | 0,006363145 |
| GAL | 2,095209135 | 0,000339293 | DSCAML1 | 2,603034024 | 0,001909552 | SPATA22 | 1,86416685 | 0,006490852 |
| FZD5 | -2,217424853 | 0,000339293 | LPAR4 | -2,840036833 | 0,002027283 | MAB21L2 | -2,344727736 | 0,006490852 |
| ERBB4 | -1,608195924 | 0,000339293 | BMP6 | 1,797677275 | 0,002067634 | PREX2 | -1,895214593 | 0,00654097 |
| JUNB | 1,651100338 | 0,000363745 | DRD2 | 1,911310104 | 0,002119699 | LRRIQ1 | -2,230304268 | 0,006747866 |
| SYTL1 | 2,38078558 | 0,000381256 | INHBA | 1,943531045 | 0,002198855 | PLXDC1 | 1,720148106 | 0,006814196 |
| APOBEC3C | 1,532547029 | 0,000381256 | HHIPL2 | -3,371136543 | 0,002252125 | CBR1 | 2,746093702 | 0,006991214 |
| SNAI2 | 1,80924925 | 0,000391447 | CRLF1 | 1,941126358 | 0,002294054 | PTPRZ1 | -2,289835098 | 0,006997138 |
| RASL10A | 1,877442599 | 0,000415202 | ADAMTSL2 | 1,594546261 | 0,002388538 | GATA3 | 1,58890351 | 0,007566377 |
| PDK4 | 2,83974063 | 0,00043132 | FZD3 | -1,5642083 | 0,00246262 | SLC6A3 | -1,725541298 | 0,007724626 |
| FBLN1 | -1,588350862 | 0,000444297 | MKI67 | -1,620940441 | 0,002506658 | EPHA4 | -1,949119966 | 0,007737852 |
| BMP3 | -1,918899291 | 0,000449801 | CXCR2 | 1,854163088 | 0,00256305 | LTBP4 | -1,510220625 | 0,008205986 |
| LAMP5 | 1,686374304 | 0,000451403 | FOSL1 | 1,871421395 | 0,002601514 | RASD1 | 1,776910188 | 0,008344487 |
| NDRG1 | 1,551820455 | 0,000451403 | ZNF385B | -1,81912287 | 0,002601514 | FAT4 | -1,540323008 | 0,008352329 |
| KCNT2 | -1,6565026 | 0,00046383 | SLC15A3 | -1,776559618 | 0,002623801 | TAS2R3 | -2,803397109 | 0,008662833 |
| PCDH18 | -1,908015374 | 0,00048586 | GPR26 | 2,554885167 | 0,002627384 | FAM228B | 1,920625724 | 0,008786462 |
| C1QL3 | 2,750006723 | 0,000496238 | PHLDA2 | 1,912747543 | 0,002627384 | CGB5 | 1,997764466 | 0,008933737 |
| SUMO4 | -1,986991477 | 0,000506112 | RRAD | 2,053971838 | 0,002728733 | DUSP2 | 1,65487497 | 0,008983737 |
| ARHGDIB | 1,850929182 | 0,000509686 | FREM2 | -1,565177808 | 0,002728733 | ROBO1 | -1,537260889 | 0,008983737 |
| PCSK6 | 1,711984469 | 0,000509686 | CAV1 | 1,914603656 | 0,002899024 | GPR183 | 2,015068932 | 0,009086429 |
| IRF1 | 1,590773748 | 0,000509686 | WNK4 | 1,997056036 | 0,002931427 | TTF1 | -1,94369392 | 0,009313694 |
| KIAA1211L | 2,76892938 | 0,000517694 | UGT3A1 | -1,686489198 | 0,003345925 | TAS2R31 | -2,897454085 | 0,009353729 |
| BBC3 | 1,535319531 | 0,000517694 | MAPRE2 | 1,610130633 | 0,003380818 | CD34 | -3,228526136 | 0,009577181 |
| ROBO2 | -2,353637478 | 0,000534263 | KDR | -1,538782171 | 0,003406179 | ZNF493 | -1,597943058 | 0,009582422 |
| MSTN | -2,419217607 | 0,000572993 | TFAP2C | 1,719741735 | 0,003623613 | OSGIN1 | 2,246915673 | 0,009631694 |
| SFRP2 | -2,547991562 | 0,00060178 | PODN | 1,685594321 | 0,003623613 | ANK1 | 1,501084248 | 0,009652588 |
| ZNF488 | 1,613702593 | 0,000668475 | ASPH | -1,680643217 | 0,003696591 | SCN3A | -1,978797967 | 0,009826333 |
| CYP26C1 | -2,60465348 | 0,000697753 | NPR1 | -1,93870055 | 0,00402619 | SAC3D1 | 1,586634217 | 0,009949286 |
| CDH6 | -1,630541842 | 0,000707553 | HES5 | -5,038583814 | 0,004080276 | TMEM168 | -2,739006335 | 0,009993603 |
| ADAMTS5 | -1,875618352 | 0,000718764 | SRGAP2 | -1,614350913 | 0,004106754 |  |  |  |
| MX1 | 3,4674981 | 0,000725331 | ISG15 | 1,862365951 | 0,004133911 |  |  |  |
| HMOX1 | 1,659405071 | 0,000725331 | BIK | 1,73069843 | 0,00451334 |  |  |  |
| HIST1H4D | -1,571617314 | 0,000814482 | CACNG7 | -1,885120997 | 0,004522138 |  |  |  |
| CDK5R2 | 1,65690867 | 0,000820304 | IRS4 | -2,193265101 | 0,004527926 |  |  |  |
| BMP5 | -1,950914501 | 0,000820304 | SLC7A8 | -1,852506781 | 0,004527926 |  |  |  |

**Table 2:**
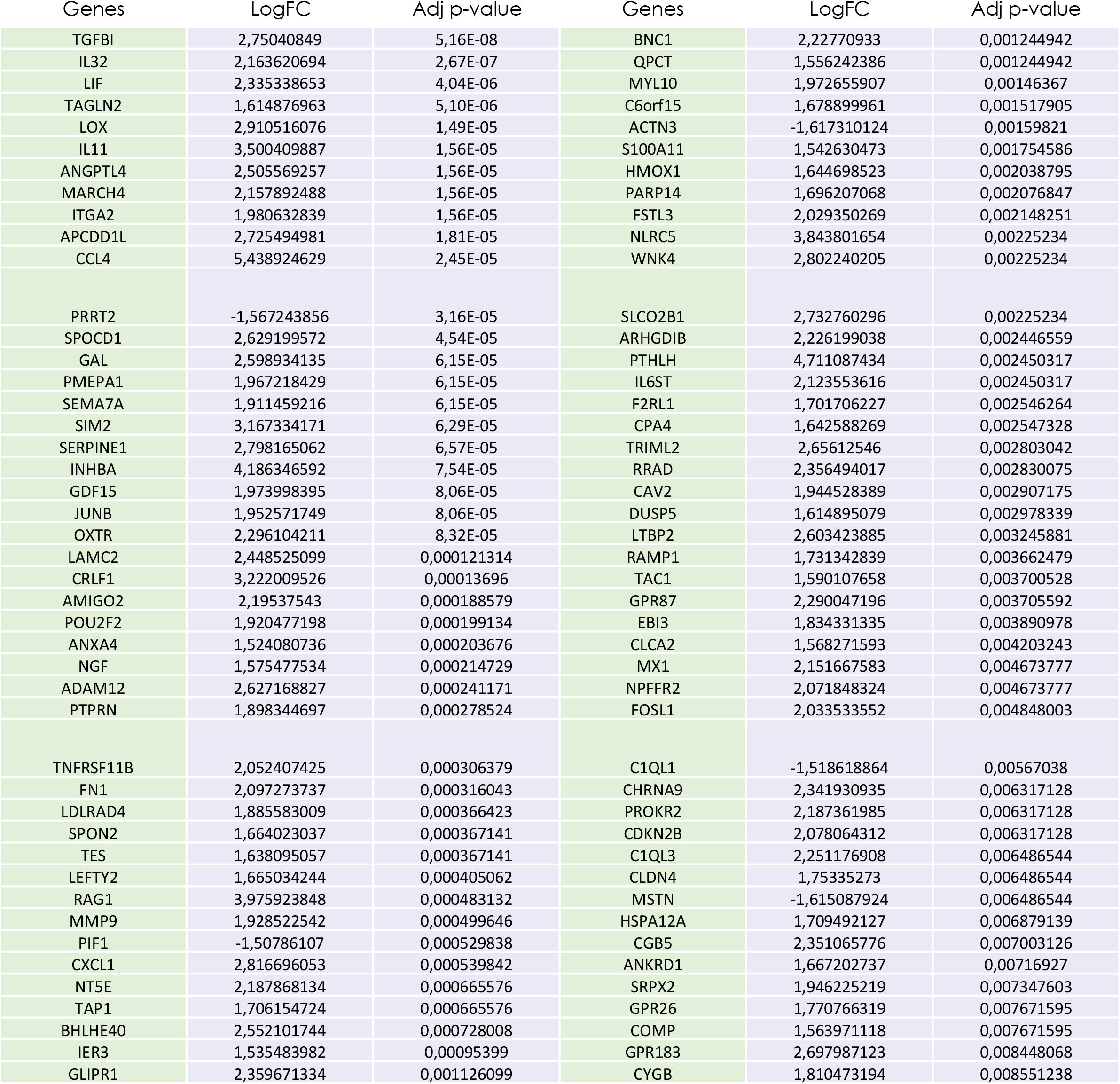
Top DEGs + *versus* – TGFB1 in DMD myotubes.

**Table 3:**
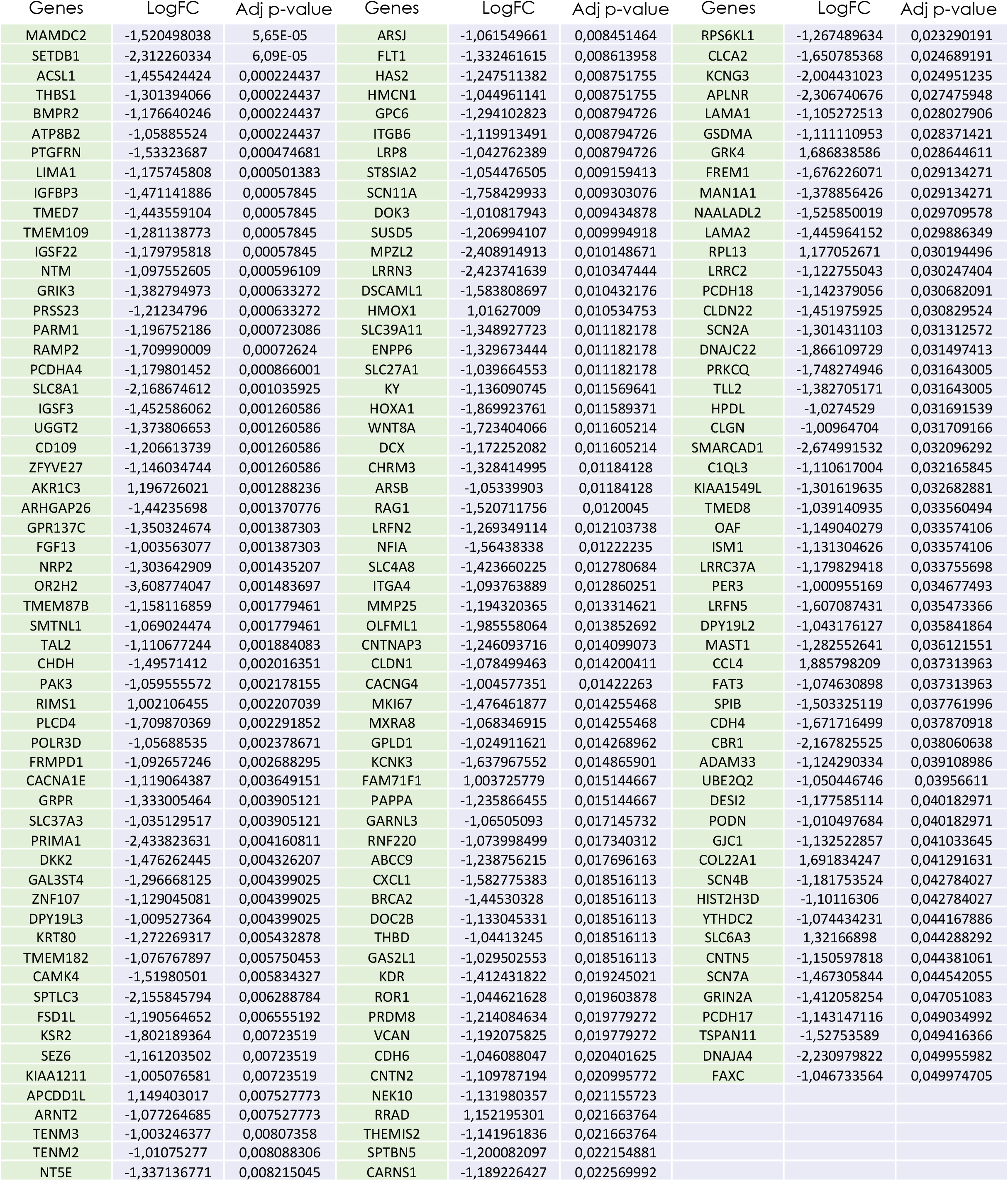
Top DEGs siSETDB1 *versus* siCTL during TGFB response in WT myotubes.

**Table 4:**
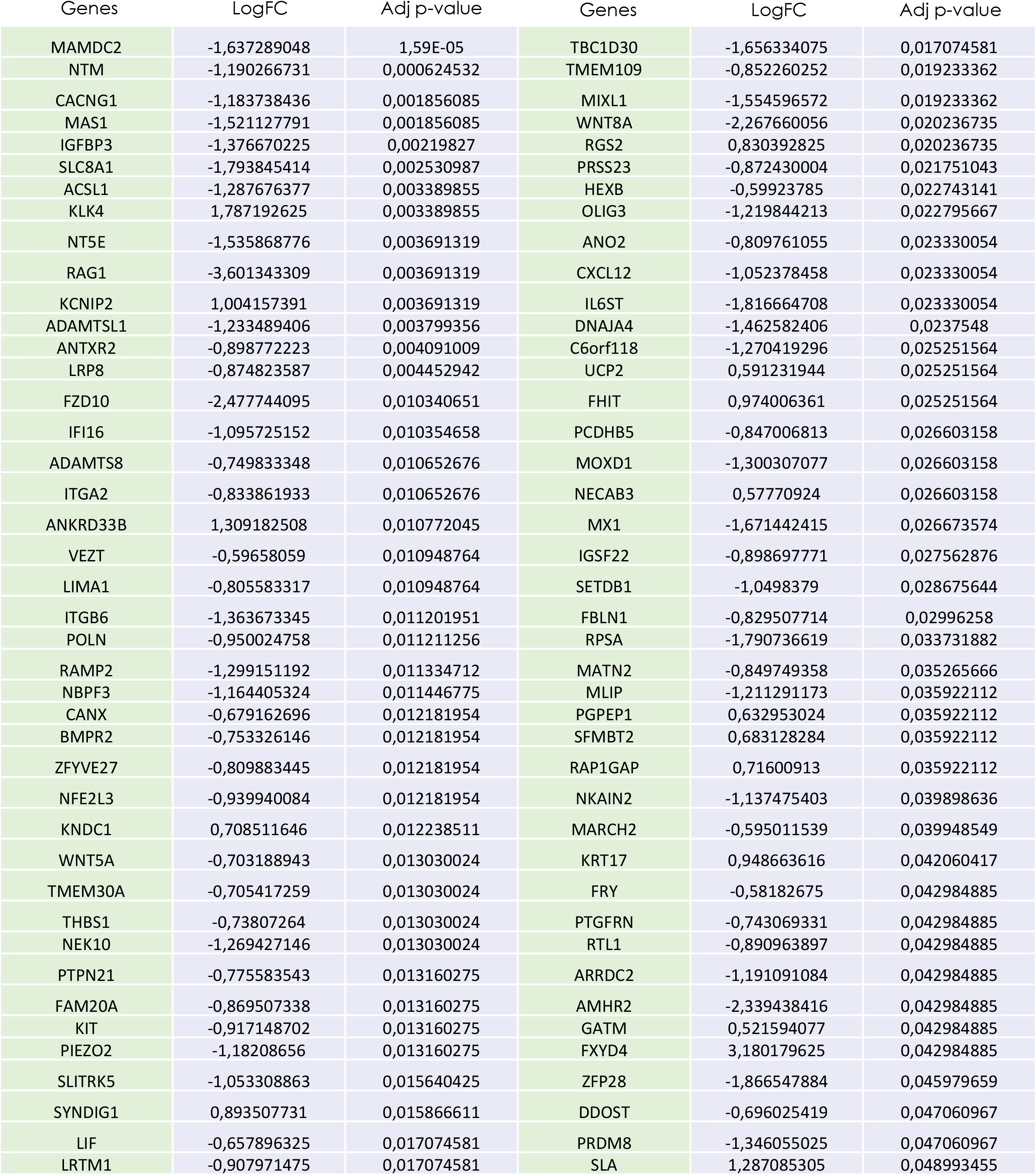
Top DEGs siSETDB1 *versus* siCTL during TGFB response in DMD myotubes.

